# Human-Specific Regulation of *CHD2* Controls the Tempo of Cortical Development through Lysosomal Homeostasis

**DOI:** 10.1101/2025.01.21.634145

**Authors:** Oliviero Leonardi, Elly Lewerissa, Davide Aprile, Tahsin Stefan Barakat, Lorenza Culotta, Ruizhi Deng, Linda Haazen, Katrin Linda, Marian Majoie, Annika Mordelt, Aurelio Ortale, Astrid Oudakker, Filippo Prazzoli, Sofia Puvogel, Nicky Scheefhals, Helenius Jurgen Schelhaas, Chantal Schoenmaker, Eline van Hughte, Hans van Bokhoven, Judith Verhoeven, Emanuele Villa, Patrycja Wypych, Alessandro Vitriolo, Cedric Boeckx, Giuseppe Testa, Nael Nadif Kasri

## Abstract

The mechanisms linking evolutionary changes in gene regulation to brain development and neurodevelopmental disease susceptibility remain poorly understood. Here, we identify a human-specific variant in an enhancer region that reduces expression of the chromatin remodeler CHD2. We investigate the variant’s functional consequences using genome editing, cross-primate induced pluripotent stem cell models, cortical organoids, single-cell transcriptomics, patient-derived cells, and neuronal network analyses. We demonstrate that CHD2 dosage bidirectionally regulates lysosomal function and autophagosome flux to set the tempo of neuronal maturation. Higher CHD2 expression, as found in ancestralized and non-human primate models, enhances lysosomal degradative capacity and accelerates dendritic and synaptic maturation. Conversely, *CHD2* haploinsufficiency yields reciprocal defects and disrupts broader neurodevelopmental transcriptional programs. Restoring lysosomal function genetically or pharmacologically rescues neuronal maturation in *CHD2*-haploinsufficient neurons, establishing lysosomal dysfunction as a causal and therapeutically tractable mechanism. These findings reveal that *CHD2* and lysosomal homeostasis constitute a critical molecular axis regulating the pace of cortical development across evolution and disease.

## INTRODUCTION

The slow-paced developmental trajectory of the human brain, often described in terms of neoteny or bradychrony, has long been proposed to contribute to the distinct cognitive and behavioral capacities of our species^1–4^. Much of our understanding of the molecular and cellular mechanisms underlying this extended developmental timeline has emerged from the functional characterization of lineage-specific genetic modifiers, such as human-specific gene duplications that have occurred during separation between us and our closest living relatives^5^. These modifiers influence several aspects of neuronal development, from the timing and complexity of cortical neurogenesis to synaptogenesis and the assembly of cortical circuits^6–8^. Beyond specific genetic changes, metabolic processes involving mitochondrial dynamics and epigenetic barriers have recently been identified as key determinants of the cell-intrinsic pace of neuronal development^9–11^.

To date, this distinct ontogeny has been characterized primarily through comparisons between modern humans and living relatives, including both closely related species (chimpanzee) and more distantly related mammals (mouse). But more subtle differences, independent of overall brain volume, could very well have existed between us and our closest extinct relatives, the Neanderthals and Denisovans. Such differences may have impacted the rate of brain maturation, as suggested in previous work^12^ based on the first draft of an extinct hominin^13^. Although endocranial volume studies establish that Neanderthal and modern human adults reached similar overall brain sizes, the distinct globular shape of *Homo sapiens*, indeed suggests a unique brain growth trajectory^14^.

Now that high-coverage genomes are available for extinct hominins^15–18^, it is possible to investigate the impact specific mutations may have had. Recent work points to differences in neocortical neurogenesis and brain metabolism^19–22^. These studies focused on mutations in protein-coding regions, which account for only a very small fraction of the genetic differences between Neanderthals/Denisovans and modern humans. Much less is known about the impact of lineage-specific mutations in regulatory regions, such as enhancers and promoters. Yet, regulatory changes have long been proposed to contribute to human evolution^23^ and are enriched in genomic regions showing signatures of positive selection^24^ or depleted of Neanderthal/Denisovan introgression^25^.

Interpreting the functional consequences of individual noncoding variants, however, remains challenging because their effects on gene regulation and neurodevelopmental phenotypes are often indirect and context-dependent. Recent work investigating species-specific regulatory differences has provided experimental validation for the importance of regulatory changes in brain growth^26,27^. Building on these advances, we previously integrated a catalog of high-frequency modern human variants with epigenomic maps, chromatin interaction datasets, and single-cell transcriptomic profiles from the developing cortex to identify regulatory elements that acquired derived variants along the modern human lineage^28,29^. Several of these candidate regulatory regions are located near dosage-sensitive neurodevelopmental genes, including *CHD8*, *SETD1A*, and *CHD2,* suggesting that regulatory evolution may have targeted pathways controlling chromatin organization and developmental timing.

*CHD2* encodes an ATP-dependent chromatin remodeler with essential functions throughout corticogenesis, including progenitor maintenance, neuronal differentiation, migration and circuit maturation^30,31^. Importantly, *CHD2* is highly dosage sensitive in humans. Haploinsufficiency causes a neurodevelopmental syndrome characterized by epilepsy, intellectual disability, and autism spectrum disorder, whereas increased *CHD2* dosage, resulting from disruption of the regulatory long noncoding RNA (IncRNA) *CHASERR*, also leads to severe developmental abnormalities^32^. These observations raise the possibility that evolutionary changes affecting CHD2 expression could have consequences for the tempo and trajectory of human cortical development.

Here, we investigate a modern human-specific regulatory variant associated with *CHD2* and examine its consequences across evolutionary and disease contexts. Using genome editing, cross-primate cellular models, cortical organoids, and patient-derived cells, we show that a single nucleotide substitution modulates CHD2 dosage and rewires neurodevelopmental transcriptional programs. We further identify lysosomal function as a major downstream effector of CHD2 dosage, linking human-specific regulatory evolution to a cellular pathway that governs cortical neuron maturation and developmental timing.

## RESULTS

### A sapiens-derived enhancer variant modulates CHD2 dosage across species

To uncover human-specific regulatory changes affecting cortical development, we revisited a set of candidate regulatory elements previously identified through the integration of high-frequency modern human variants with epigenomic, chromatin interaction, and developmental transcriptomic datasets from the fetal cortex^28,29^. To determine which variants exert the most substantial functional influence, we applied BRAIN-MAGNET, a convolutional neural network designed to estimate how noncoding variants affect gene-level transcriptional variance^33^. BRAIN-MAGNET prioritized a high-frequency derived variant located within an enhancer approximately 77 kb upstream of *CHD2*, ranking it within the top 5% of all candidates identified in our screen (Figure 1A). While nearly fixed in present-day humans, the sapiens-derived (hereafter derived) C allele is absent from Neanderthal and Denisovan genomes, which retain the ancestral T allele shared with chimpanzees, bonobos, and other primates (Figure 1B). Hi-C data from fetal cortex revealed a physical interaction between this enhancer and the promoter of *CHD2*. (Figure S1A). The enhancer is also enriched for chromatin marks associated with active regulatory elements, including H3K4me1, H3K4me3, H3K9ac, and H3K27ac, in both pluripotent stem cells and neuronal tissues (Figures S1B). BRAIN-MAGNET nucleotide-level contribution scores further indicates that the variant itself lies within one of the sequence positions predicted to contribute strongly to enhancer activity, suggesting a direct effect on transcription factor binding (Figure 1C).

**Figure 1.**
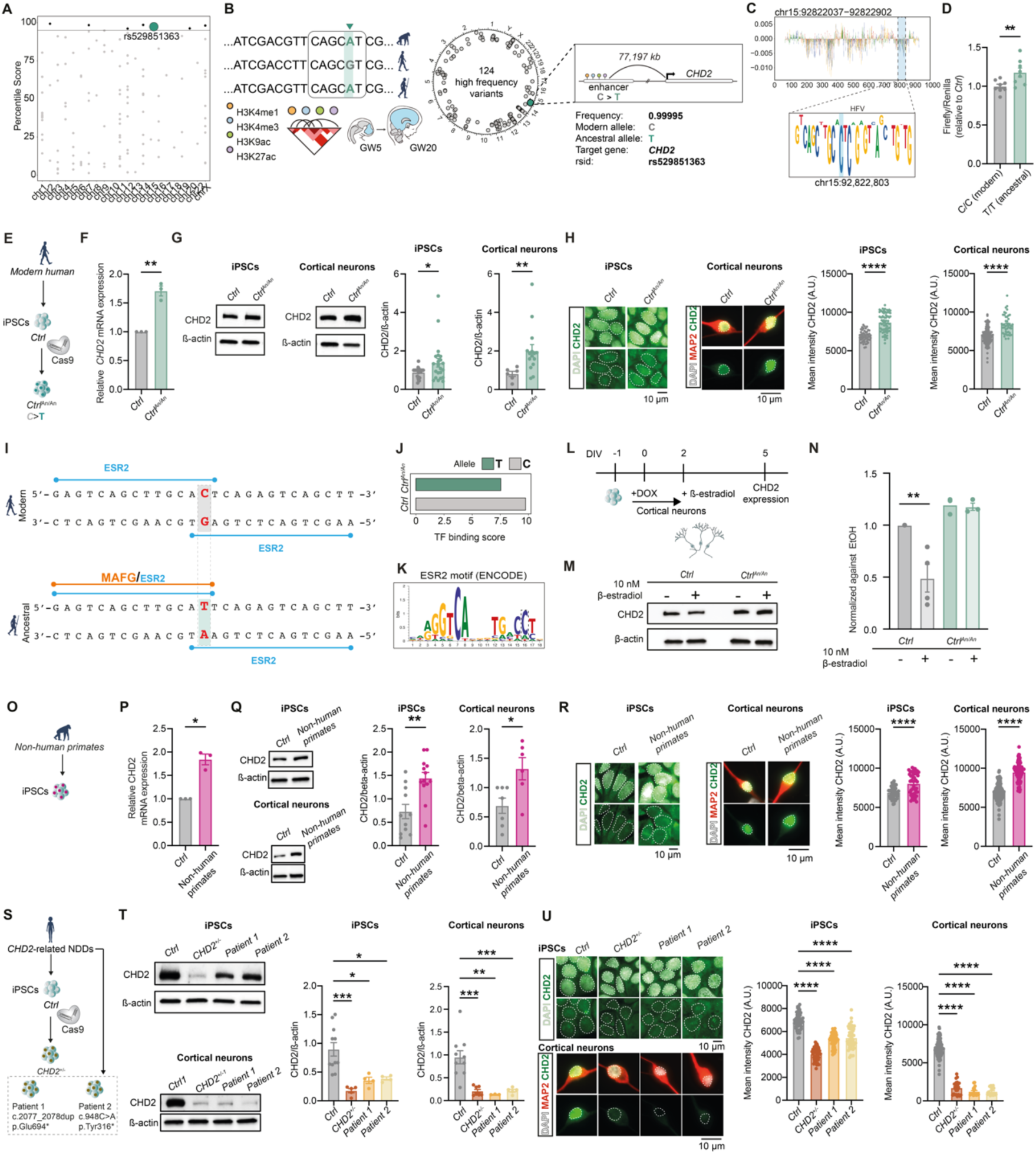
CHD2 dosage is modulated by a contemporary-human enhancer variant and altered in CHD2-related evolutionary and neurodevelopmental disorder models. (A) BRAIN-MAGNET percentile scores for the 124 prioritized high-frequency variants, with the *CHD2*-linked rs529851363 variant highlighted. (B) Schematic of the BRAIN-MAGNET-based prioritization strategy. Modern-human high-frequency variants overlapping regulatory regions annotated from early cortical developmental epigenomic datasets were screened, identifying 124 candidate variants. The highlighted variant, rs529851363, maps to an enhancer located approximately 77 kb upstream of *CHD2*; the modern-human allele is C and the ancestral allele is T. (C) BRAIN-MAGNET nucleotide-level attribution profile across the *CHD2* enhancer, illustrating the contribution of individual nucleotides to the predicted transcriptional effect of the enhancer. The sequence logo highlights the region centered on rs529851363. (D) Luciferase reporter assay in neural stem cells comparing enhancer activity driven by the modern C allele or the ancestral T allele upstream of a luciferase reporter. Firefly luciferase activity was normalized to Renilla and is shown relative to modern allele. Each dot represents an independent transfection, data are pooled from two independent experiments (n = 8 per condition). (E) Schematic of CRISPR-Cas9 editing of human *Ctrl* iPS cells to generate isogenic *Ctrl^An/An^*lines carrying the ancestral T allele at the CHD2 enhancer. (F) qPCR quantification of *CHD2* mRNA expression in *Ctrl* and *Ctrl^An/An^* iPS cells. Each dot represents one independent qPCR (n = 3). (G) Representative western blots and quantification of CHD2 protein levels in *Ctrl* and *Ctrl^An/An^* iPS cells and DIV 5 cortical neurons. β-actin was used as loading control. Each dot represents the quantification of one independent western blot; data are pooled from five independent western blot experiments (n = 24 for *Ctrl* iPS cells, n = 22 for *Ctrl^An/An^* iPS cells; n = 6 for *Ctrl* neurons, n = 15 for *Ctrl^An/An^* neurons). (H) Representative immunocytochemistry images and quantification of CHD2 intensity in *Ctrl* and *Ctrl^An/An^* iPS cells and DIV 5 cortical neurons. Cortical neurons were co-stained for MAP2, and nuclei were counterstained with DAPI. Scale bars, 10 μm. Each dot represents the quantification of one cell; data are pooled from three independent experiments (n = 50 for *Ctrl* iPS cells, n = 73 for *Ctrl^An/An^* iPS cells; n = 133 for *Ctrl* neurons, n = 45 for *Ctrl^An/An^* neurons). (I) Predicted transcription factor binding site architecture around rs529851363 in the modern and ancestral enhancer sequences. The modern C allele is predicted to increase the binding affinity of ESR2, whereas the ancestral T allele creates an overlapping MAFG/ESR2 configuration. (J) ENCODE ESR2 motif logo used for predictions. (K) Predicted ESR2 transcription factor binding score for the modern C and ancestral T alleles. (L) Experimental scheme for 17β-estradiol treatment in *NGN2*-induced cortical neurons. Neurons were treated with 10 nM 17β-estradiol or vehicle at the indicated timepoints, and collected for CHD2 protein analysis at DIV 5. (M) Representative western blot of CHD2 protein levels in *Ctrl* and *Ctrl^An/An^* neurons treated with 10 nM 17β-estradiol or vehicle (ethanol). Β-actin was used as loading control. (N) Quantification of CHD2 protein levels shown in (M), normalized to the corresponding vehicle-treated condition. Each dot represents the quantification of one biological experiment; data are pooled from three independent experiments. (n = 4 for *Ctrl* and *Ctrl +*10 nM 17β-estradiol, n = 3 for *Ctrl^An/An^* and *Ctrl^An/An^* +10 nM 17β-estradiol). (O) Schematic of the comparison between human control and non-human primate iPS cells. (P) qPCR quantification of *CHD2* mRNA expression in human *Ctrl* and non-human primate iPS cells. Each dot represents one independent qPCR (n = 3 for *Ctrl*, n = 3 for non-human primates [n = 2 for chimpanzee, n = 1 for bonobo]). (Q) Representative Western blots and quantification of CHD2 protein levels in human *Ctrl* and non-human primate iPS cells and DIV 5 cortical neurons. β-actin was used as loading control. Each dot represents the quantification of one independent Western blot; data are pooled from three independent western blot experiments (n = 6 for *Ctrl*, n = 6 for non-human primates [n= 3 for bonobo, n = 3 for chimpanzee]). (R) Representative immunocytochemistry images and quantification of CHD2 intensity in human *Ctrl* and non-human primate iPS cells and DIV 5 cortical neurons. Cortical neurons were co-stained for MAP2, and nuclei were counterstained with DAPI. Scale bars, 10 μm. Each dot represents the quantification of one independent cell; data are pooled from three independent experiments (n = 50 for *Ctrl* iPS cells, n = 50 for non-human primes [n = 25 for bonobo, n = 25 for chimpanzee], n = 133 for *Ctrl* neurons, n = 97 for non-human primates [chimpanzee]). (S) Schematic of CHD2-related neurodevelopmental disorder models used in this study, including CRISPR-Cas9-generated isogenic *CHD2^+/−^* iPS cells and two independent patient-derived iPS cell lines (*patient* 1 and *patient* 2) carrying pathogenic CHD2 variants. (T) Representative western blots and quantification of CHD2 protein levels in *Ctrl*, *CHD2^+/−^*, *patient 1*, and *patient 2* iPS cells and DIV 5 cortical neurons. β-actin was used as loading control. Each dot represents the quantification of one independent western blot; data are pooled from three independent western blot experiments (n = 11 for *Ctrl* iPS cells, n = 6 for *CHD2^+/−^* iPS cells, n = 4 for *patient 1* iPS cells, n = 4 for *patient 2* iPS cells; n = 10 for *Ctrl* neurons, n = 7 for *CHD2^+/−^* neurons, n = 3 for *patient 1* neurons, n = 5 for *patient 2* neurons). (U) Representative immunocytochemistry images and quantification of CHD2 intensity in *Ctrl*, *CHD2^+/−^*, *patient 1*, and *patient 2* iPS cells and DIV 5 cortical neurons. Cortical neurons were co-stained for MAP2, and nuclei were counterstained with DAPI. Scale bars, 10 μm. Each dot represents the quantification of one independent cell; data are pooled from three independent experiments (n = 50 for *Ctrl* iPS cells, n = 50 for *CHD2^+/−^* iPS cells, n = 50 for *patient 1* iPS cells, n = 50 for *patient 2* iPS cells; n = 133 for *Ctrl* neurons, n = 24 for *CHD2^+/−^* neurons, n = 18 for *patient 1* neurons, n = 21 for *patient 2* neurons). Data represent means ±SEM. *p<0.05, **p<0.01, ***p<0.01, ****p<0.0001, one-way ANOVA with post hoc Bonferroni correction, and unpaired two-tailed student t-test.

To test the consequences of this variant on gene expression, we performed a luciferase reporter assay in neural stem cells. The derived C allele exhibited significantly lower enhancer activity than the ancestral T allele present in Neanderthals, Denisovans, and great apes (Figure 1D). To determine whether this single nucleotide change is sufficient to alter endogenous CHD2 expression, we used CRISPR–Cas9 to revert the derived allele to the ancestral state in two independent human control iPS cell lines, generating isogenic ancestralized (*Ctrl^An/An^*) lines (Figure 1E). Precise editing was confirmed by Sanger sequencing (Figure S2A-B). Potential off-target events at genomic sites carrying up to three mismatches to the guide RNA were excluded by targeted sequencing (Figure S2C-D), and genomic integrity was verified by digital droplet PCR (Figure S2E) and whole-genome sequencing (Figure S2F). Consistent with the reporter assays, Ctrl*^An/An^*iPS cells displayed increased *CHD2* mRNA expression relative to isogenic controls (Figure 1F), demonstrating that this single nucleotide substitution is sufficient to modulate transcription from the endogenous locus. Elevated CHD2 expression was also detected at the protein level in both iPS cells and iPS cell-derived cortical neurons by western blot analysis (Figure 1G). Immunocytochemistry confirmed these differences in both cell types (Figure 1H). Together, these findings indicate that the derived enhancer allele reduces CHD2 expression in human pluripotent stem cells and cortical neurons.

To investigate the mechanism underlying this regulatory effect, we compared predicted transcription factor binding site architectures between the ancestral and derived enhancer sequences. Analysis of a ± 20 bp window surrounding the variant identified all transcription factors with high-confidence binding sites in either allele (Figure 1I, Figure S3A-B). In the ancestral sequence, the variant overlaps predicted binding sites for MAFG and ESR2, creating potential competition between these factors (binding affinity scores: MAFG = 8.25; ESR2 = 7.55). The derived T>C substitution disrupts the MAFG binding site while increasing the predicted affinity of ESR2 to 9.75 (Figure 1J). Quantitative motif analysis confirmed that the derived C allele matches more closely to the canonical ESR2-binding motif defined by ENCODE than the ancestral allele (Figure 1K). Consistent with these predictions, ESR ChIP-seq datasets from ReMap^34^ revealed recurrent binding peaks centered on the variant across multiple human cell types (Figure S4), supporting the interpretation that this region functions as an estrogen-responsive regulatory element. To functionally test these predictions, cortical neurons derived from control (*Ctrl*) and *Ctrl^An/An^* iPS cells were treated with 17β-estradiol or vehicle, and CHD2 protein levels were quantified by western blot (Figure 1L). In *Ctrl* neurons carrying the derived allele, estradiol treatment induced an approximately two-fold reduction in CHD2 protein levels. In contrast, *Ctrl^An/An^* neurons carrying the ancestral allele showed no significant response to estradiol exposure (Figure 1M-N). These findings indicate that the derived human allele increases the sensitivity of CHD2 regulation to estrogen signaling.

Because the derived enhancer variant reduced CHD2 expression in human cells, we next examined whether this effect was consistent in a broader evolutionary context. To this end, we compared human (*Homo sapiens*), chimpanzee (*Pan troglodytes*), and bonobo (*Pan paniscus*) cellular models (Figure 1O). Consistent with the ancestralized human lines, both chimpanzee and bonobo iPS cells exhibited higher *CHD2* mRNA expression than human *Ctrl* iPS cells (Figure 1P). Increased CHD2 protein levels were likewise observed in both iPS cells and cortical neurons derived from non-human primates (Figure 1Q-R). Although these broader cross-species comparisons may also reflect the cumulative effects of additional regulatory differences beyond the enhancer variant studied here, the direction of change is consistent with higher ancestral CHD2 expression and reduced expression along the modern human lineage.

Finally, to directly examine the consequences of reduced CHD2 dosage, we generated complementary disease-relevant models. These included two patient-derived iPS cell lines carrying pathogenic *CHD2* mutations (*Patient 1* and *Patient 2*, (Table S1) and isogenic heterozygous *CHD2* loss-of-function lines generated by CRISPR–Cas9 in two independent genetic backgrounds (*CHD2^+/-^*) (Figure 1S, Figure S5). Western blotting and immunocytochemistry demonstrated a marked reduction in CHD2 protein levels in both *CHD2^+/-^*and patient-derived cells relative to their respective controls, in both iPSCs and cortical neurons (Figure 1T-U). Collectively, these findings identify CHD2 as a dosage-sensitive regulator whose expression varies across evolutionary and disease contexts: elevated in ancestralized human and non-human primate cells while reduced in neurodevelopmental disorders caused by *CHD2* haploinsufficiency.

### CHD2 dosage modulates disease-associated and autolysosomal transcriptional programs

Building on these species- and allele-specific differences in CHD2 expression, we next examined the downstream transcriptional consequences during neuronal development. We performed bulk RNA-sequencing on DIV 5 cortical neurons and compared each genotype with its isogenic control. In these early cortical neurons, *CHD2^+/-^* cells displayed 2,273 differentially expressed genes (DEGs), while *Ctrl^An/An^* neurons exhibited 738 DEGs (Figure 2A). Direct comparison of differential expression profiles revealed 921 genes uniquely altered in *CHD2^+/-^* neurons, 124 uniquely altered in *Ctrl^An/An^* neurons, and 118 genes shared between the two conditions (Figure 2B). Notably, most shared genes were regulated in opposite directions, indicating that increased and decreased CHD2 dosage can converge on overlapping transcriptional targets while exerting reciprocal effects on their expression (Figure 2B).

**Figure 2.**
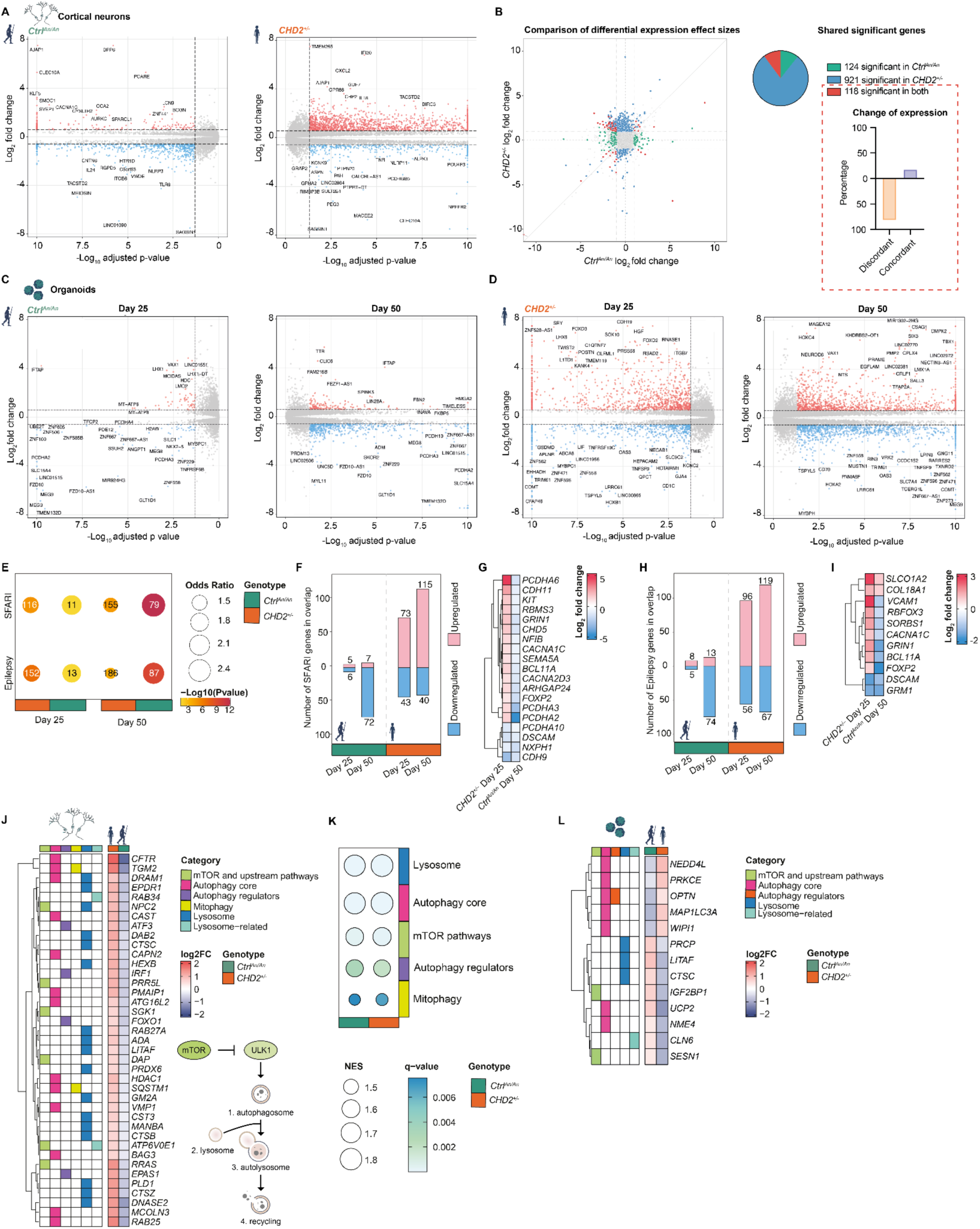
CHD2 dosage remodels disease-associated and autolysosomal transcriptional programs. (A) Volcano plots showing differential gene expression in DIV 5 cortical neurons comparing *Ctrl^An/An^*and *CHD2^+/−^* neurons with their corresponding isogenic controls. Red and blue dots indicate significantly upregulated and downregulated genes, respectively; gray dots indicate genes not passing the differential expression threshold. Selected genes are labeled. DEGs were defined as FC ≥ |1.5| and FDR ≤ 0.05; n = 3. (B) Scatter plot comparing differential expression effect sizes between *Ctrl^An/An^* and *CHD2^+/−^* cortical neurons. Each dot represents one gene, green colored according to whether it is significantly altered only in *Ctrl^An/A^* neurons, blue colored only in *CHD2^+/−^*neurons, or in both comparisons (red colored). The pie chart summarizes the number of significant genes in each category, and the bar plot shows the percentage of shared DEGs changing in concordant or discordant directions between genotypes. (C-D) Volcano plots showing differential gene expression in *Ctrl^An/An^* or *CHD2^+/−^* cortical organoids at day 25 and day 50 compared with isogenic control organoids collected at the same developmental stages. DEGs were defined as FC ≥ |1.5| and FDR ≤ 0.05; n = 3. (E) Dot plot showing the overlap between organoid DEGs and curated SFARI or epilepsy-associated gene sets across genotypes and developmental stages. Dot size indicates the odds ratio, color indicates −log10(p value), and numbers inside dots indicate the number of overlapping genes. Overlap significance was assessed using Fisher’s exact test, with all tested genes used as background. (F) Number and direction of SFARI genes overlapping with organoid DEGs. Bars indicate upregulated and downregulated genes, separated by genotype and developmental stage. (G) Heatmap showing log2 fold changes for selected SFARI genes differentially expressed in *CHD2^+/−^* day 25 organoids and *Ctrl^An/An^* day 50 organoids. (H) Number and direction of epilepsy-associated genes overlapping with organoid DEGs. Bars indicate upregulated and downregulated genes, separated by genotype and developmental stage. (I) Heatmap showing log2 fold changes for selected epilepsy-associated genes differentially expressed in *CHD2^+/−^* day 25 organoids and *Ctrl^An/An^* day 50 organoids. (J) Heatmap of autophagy-lysosome pathway-related DEGs in cortical neurons. The left annotation matrix indicates pathway membership, including mTOR and upstream pathways, autophagy core genes, autophagy regulators, mitophagy, lysosome-associated genes, and lysosome-related genes. The right heatmap shows log2 fold changes in *Ctrl^An/An^* and *CHD2^+/−^* neurons relative to their corresponding controls. Inset shows a schematic overview of the autophagy-lysosome pathway, illustrating autophagosome formation downstream of mTOR-ULK1 signaling, fusion with lysosomes to generate autolysosomes, and subsequent cargo degradation and recycling. (K) Gene set enrichment analysis (GSEA) of autophagy-lysosome pathway categories in *Ctrl^An/An^* and *CHD2^+/−^* cortical neurons at DIV 5. Dot size indicates the normalized enrichment score (NES), and color indicates the q value. (L) Heatmap of autophagy-lysosome pathway-related DEGs in cortical organoids across genotype and developmental stage. The annotation matrix indicates pathway membership, including mTOR and upstream pathways, autophagy core genes, autophagy regulators, lysosome-associated genes, and lysosome-related genes. The expression heatmap shows log2 fold changes relative to the corresponding control condition.

To determine how these dosage-dependent effects evolve during cortical development, we profiled cortical organoids at days 25 and 50, corresponding to distinct stages of early corticogenesis. Both *Ctrl^An/An^*and *CHD2^+/−^* organoids displayed developmental stage-dependent transcriptional changes, although the magnitude of dysregulation was consistently greater in *CHD2*^+/−^ organoids at both time points (Figure 2C-D). Given the established association of *CHD2*-related disorders with autism spectrum disorder (ASD) and epileptic encephalopathy, we examined whether differentially expressed genes were enriched for disease-associated gene sets. Intersecting organoid DEGs with curated SFARI and epilepsy-associated gene lists revealed significant enrichment in both models, although the effect was markedly stronger and more persistent in *CHD2*^+/−^ organoids (Figure 2E). In *CHD2*^+/−^ organoids, 116 and 155 SFARI genes overlapped with DEGs at days 25 and 50, respectively, while 152 and 186 epilepsy-associated genes were differentially expressed at the corresponding stages. *Ctrl^An/An^* organoids also showed significant overlap, particularly at day 50, when 79 SFARI genes and 87 epilepsy-associated genes were differentially expressed (Figure 2E). Interestingly, the direction of these transcriptional changes was largely opposite between the two models. The ASD- and epilepsy- associated genes that were predominantly upregulated in *CHD2*^+/−^ organoids at day 25 were downregulated in *Ctrl^An/An^* organoids, particularly at day 50 (Figure 2F,H). Extending this observation, CHD2 dosage reciprocally regulates a core set of disease-relevant genes, including several dosage- sensitive regulators of neuronal identity, adhesion, excitability, and circuit formation, including *GRIN1, CACNA1C, BCL11A, FOXP2, DSCAM*, protocadherins, and cadherins (Figure 2G,I). Together, these findings indicate that the evolutionary increase in CHD2 dosage and disease-associated *CHD2* haploinsufficiency converge on overlapping neurodevelopmental pathways, while driving opposite transcriptional outcomes across shared gene networks.

Notably, reciprocal transcriptional changes also extended to genes involved in mTOR signaling, core autophagy machinery, autophagy regulators, lysosomes, and lysosome-associated pathways (Figure 2J). Recent studies have linked species-specific differences in developmental timing to metabolic and proteostatic pathways^35,36^, including those controlled by autophagy and lysosomal function during neural differentiation. Gene set enrichment analysis (GSEA) identified significant enrichment of *autophagy*- and *lysosome*-related pathways in both *Ctrl^An/An^* and *CHD2^+/−^* neurons (Figure 2K), suggesting that the autophagy-lysosome pathway represents a common downstream target of CHD2 dosage variation. This transcriptional signature was recapitulated in cortical organoids, where genes involved in mTOR signaling, autophagy regulation, autophagosome formation, lysosomes, and lysosome-associated function were differentially regulated across genotypes and developmental stages (Figure 2L). While enhancer ancestralization and *CHD2* haploinsufficiency drive reciprocal transcriptional changes, both perturbations converge on the autophagy-lysosome pathway, implicating lysosome-dependent proteostasis as a shared downstream consequence of altered CHD2 dosage.

### CHD2 dosage tunes lysosomal degradative capacity

The reciprocal regulation of autophagy- and lysosome-associated gene programs in *Ctrl^An/An^* and *CHD2^+/-^* cortical neurons suggested that CHD2 dosage influences the functional state of the autophagy- lysosome pathway. We therefore examined multiple functional aspects of this pathway, including lysosomal acidification, proteolytic activity, lysosomal dynamics, and autophagosome flux across evolutionary and disease-associated CHD2 models. Using LysoSensor, a pH-sensitive dye that fluoresces more brightly in acidic environments (Figure 3A), we observed significantly increased lysosomal acidification in *Ctrl^An/An^* neurons relative to *Ctrl*, while *CHD2^+/-^* neurons showed reduced lysosomal acidity (Figure 3A). Consistent with the elevated CHD2 expression observed in non-human primate neurons, bonobo and chimpanzee neurons similarly exhibited enhanced lysosomal acidification compared to human controls (Figure 3A). Because lysosomal acidification is required for proteolytic activity, we next examined cathepsin B processing by western blot (Figure 3B). *Ctrl^An/An^* neurons exhibited increased maturation of cathepsin B, whereas *CHD2^+/-^* and patient-derived neurons accumulated immature pro-cathepsin B, resulting in a reduced cathepsin B maturation index (Figure 3C). Consistent with these findings, RNA-sequencing revealed reciprocal regulation of multiple cathepsin family members across CHD2 dosage models (Figure S6A), supporting a bidirectional effect of CHD2 dosage on lysosomal protease regulation.

**Figure 3.**
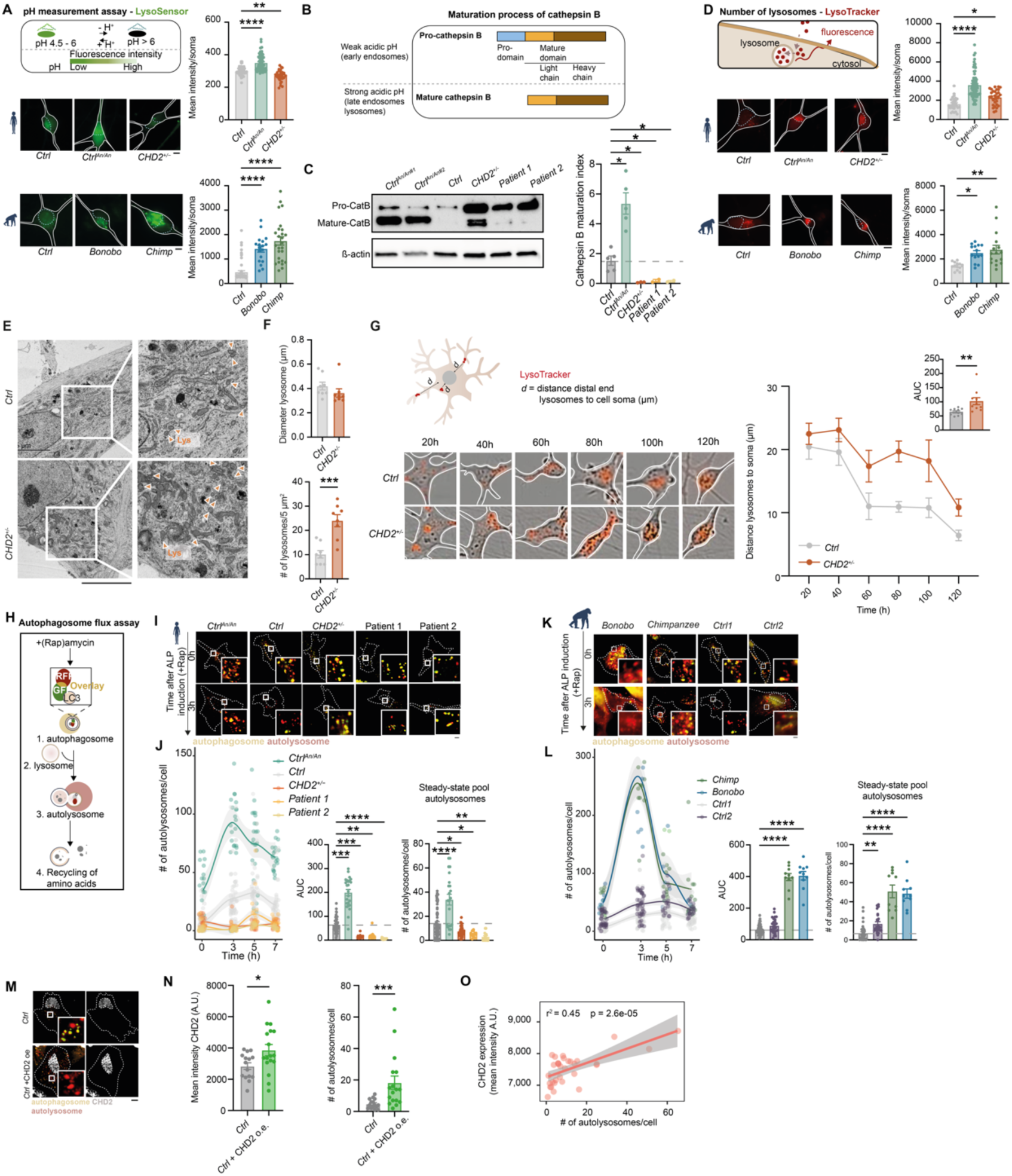
CHD2 dosage controls lysosomal degradative capacity and autophagosome flux. (A) Schematic of the LysoSensor green assay (top), where pH > 6 reduces fluorescence. Representative images of LysoSensor in cortical neurons derived from *Ctrl^An/An^*, *Ctrl*, *CHD2^+/-^* and non-human primates are shown (bottom). Dashed light blue circles indicate the position of the DAPI-stained nucleus. Quantification of LysoSensor intensity in neurons (n = 48 for *Ctrl*, n = 68 for *Ctrl^An/An^*, n = 28 for *CHD2^+/-^*, n = 18 for *bonobo* and n = 26 for *chimpanzee*) (right). (B) Schematic overview of cathepsin B maturation. Pro-cathepsin B is proteolytically processed within acidic lysosomes to generate mature cathepsin B. (C) Representative western blot and quantification of the cathepsin B maturation index in *Ctrl*, *Ctrl^An/An^*, and *CHD2*-haploinsuffient (*CHD2^+/−^*and patient 1 and patient 2) -derived cortical neurons. The maturation index was calculated as the ratio of mature cathepsin B to total cathepsin B protein. β-actin served as a loading control. Each dot represents the quantification of one independent Western blot; data are pooled from three independent western blot experiments (n = 5 for *Ctrl*, n = 5 for *Ctrl^An/An^*, n = 3 for *CHD2^+/-^*, n = 3 for *patient 1* and n = 3 for *patient 2*). (D) (top) Schematic of LysoTracker, a dye for endolysosomal staining. (bottom) Representative LysoTracker images in neurons derived from *Ctrl^An/An^*, *Ctrl*, *CHD2^+/-^* and non-human primates. Dashed light blue circles indicate the position of the DAPI-stained nucleus. (right) Quantification of LysoTracker intensity in cortical neurons (n = 28 for *Ctrl*, n = 69 for *Ctrl^An/An^*, n = 30 for *CHD2^+/-^*, n = 11 for *bonobo* and n = 16 for *chimpanzee*). (E) Transmission electron microscopy images showing lysosomes (indicated by the orange arrows) in *Ctrl* and *CHD2^+/-^*-derived neurons. (F) (top) Quantification of the number of lysosomes per 5 mm^2^ and (bottom) quantification of the diameter of individual lysosomes (mm) in *Ctrl* compared to *CHD2^+/-^*-derived neurons (n = 8 for *Ctrl* and n = 8 for *CHD2^+/-^*). (G) (top) Schematic illustrating the measurement of the distance between distal lysosomes and the neuronal soma. Representative LysoTracker images showing lysosome localization over time in *Ctrl* and *CHD2^+/-^*neurons. (right) Quantification of the average distance from distal lysosomes to the soma over time in *Ctrl* and *CHD2^+/-^* neurons. The inset shows the area under the curve (AUC) for lysosome-to-soma distance measurements (n = 10 for *Ctrl* and n = 10 for *CHD2^+/-^*). (H) Schematic representation of autophagosome flux assay. (I) Representative images of autophagosomes (yellow) and autolysosomes (red) at steady-state (0h) and after 200 nM rapamycin (Rap) (3h) treatment in iPS cells (circled by white stripes) derived from *Ctrl^An/An^*, *Ctrl*, and *CHD2*-haploinsufficient (*CHD2^+/-^,* patient 1, and patient 2). (J) (left) Quantification of autolysosomes per cell at 0h and after 200 nM rapamycin (Rap) treatment. (middle) Area under the curve (AUC) for the number of autolysosomes upon rapamycin induction over time. (right) Steady-state pool of autolysosomes per cell from *Ctrl^An/An^*, *Ctrl, CHD2^+/-^*, patient 1 and patient 2. (n = 22 for *Ctrl^An/An^*, n = 30 for *Ctrl*, n = 18 for *CHD2^+/-^*, n = 15 for patient 1, n = 15 for patient 2. (K) Representative images of autophagosome flux assay in *chimpanzee*- and *bonobo* compared to two independent human controls (*Ctrl1* and *Ctrl2)*. (L) (left) Quantification of autolysosomes per cell at 0h and after 200 nM rapamycin (Rap) treatment in non-human primates. (middle) Area under the curve (AUC) for the number of autolysosomes upon rapamycin induction over time. (right) Steady-state pool of autolysosomes per cell derived from *chimpanzee*, *bonobo* and human controls (*Ctrl1* and *Ctrl2*) (n = 56 for *Ctrl1*, n = 42 for *Ctrl2*, n = 10 for *bono* and n = 11 for *chimpanzee*). (M) Representative images of autophagosomes (yellow) and autolysosomes (red) at steady-state in *Ctrl* and *Ctrl* +CHD2oe iPS cells with ectopic expression of CHD2. CHD2 expression is shown in grey. (N) (left) Quantification of CHD2 expression levels and (right) the number of autolysosomes in *Ctrl* and *Ctrl* cells with ectopic expression of CHD2 (n = 16 for *Ctrl*, n = 15 for *Ctrl* +CHD2oe). (O) Linear regression analysis showing correlation between CHD2 and number of autolysosomes. Scale bar represents 10 μm. Data represent means ±SEM. *p<0.05, **p<0.01, ****p<0.0001, one-way ANOVA with post hoc Bonferroni correction, and unpaired two-tailed student t-test.

To determine whether impaired lysosomal function was accompanied by changes in endolysosomal organization, we next examined the abundance and intracellular positioning of acidic endolysosomal compartments. LysoTracker labeling revealed an increased LysoTracker-positive compartment in *Ctrl^An/An^* neurons compared with *Ctrl* neurons (Figure 3D). A similar increase was observed in bonobo and chimpanzee neurons (Figure 3D), consistent with enhanced lysosomal activity across species with higher CHD2 dosage. Surprisingly, *CHD2^+/-^* neurons also exhibited an increased LysoTracker-positive compartment despite reduced lysosomal acidification and proteolytic activity (Figure 3D). Transmission electron microscopy analysis further corroborated a significant accumulation of lysosomes in *CHD2^+/-^* neurons compared with *Ctrl* neurons (Figure 3E-F). This increase occurred without alterations in lysosomal diameter (Figure 3F), suggesting that *CHD2* haploinsufficiency leads to an expansion in lysosome number rather than changes in lysosomal morphology. Notably, the abundance of RAB7A-positive late endosomes did not differ between genotypes (Figure S6B), indicating that the ultrastructural accumulation is unlikely to be explained by an expansion of the late endolysosomal compartment. We further investigated lysosomal trafficking by longitudinal live-cell imaging of LysoTracker-labeled organelles (Figure 3G). As mature degradative lysosomes are enriched within the soma, lysosomes in *Ctrl* neurons progressively redistributed from distal neurites toward the cell body during development (Figure 3G). This developmental redistribution was delayed in *CHD2^+/-^* neurons, resulting in increased soma-to-lysosome distances during neuronal differentiation (Figure 3G).

Because lysosomal acidification and cathepsin activation are critical determinants of autophagy-lysosome pathway function, we next assessed autophagosome flux^37^ using the tandem mCherry-GFP-LC3 reporter, which distinguishes autophagosomes from autolysosomes. In autophagosomes, both tags emit fluorescence, but the acidic pH in autolysosomes quenches the GFP signal (Figure 3H-I,K). The number of autolysosomes was quantified to determine the autophagosome flux, *J*, which represents the time-dependent change in the total autolysosomal pool size before and after activation of the autophagy-lysosome pathway (e.g., the formation of autolysosomes/cell/h) (Figure S6C). Additionally, we calculated the transition time, *tnAL*, which reflects the time required for the autophagy-lysosome pathway to turn over its autolysosomal pool (Figure S6C). In *Ctrl* cells, approximately 10 autolysosomes were formed per hour upon activation of the autophagy-lysosome pathway (Figure 3I-J, Figure S6C). By contrast, *Ctrl^An/An^* cells showed a significantly higher basal (steady-state) autolysosomal count compared to *Ctrl* lines (Figure 3J, Figure S6C). Upon induction of the pathway, *Ctrl^An/An^* cells exhibited a substantial increase in autolysosome formation in response to rapamycin treatment (±20 autolysosomes/cell/h). Conversely, *CHD2*-haploinsuffient cells (*CHD2^+/-^,* patient 1, patient 2) exhibited a markedly reduced basal autolysosome count and significantly lower turnover rate compared with *Ctrl* (±2 autolysosomes/cell/h) (Figure 3I-J, Figure S6C). To determine whether the reduced autophagosome flux observed in *CHD2^+/-^* cells resulted from impaired autophagy induction or defective lysosomal degradation, we examined key regulators of autophagy initiation. Following rapamycin treatment, both *Ctrl* and *CHD2^+/-^* neurons exhibited comparable reductions in mTOR (Ser2448) and ULK1 (Ser757) phosphorylation, indicating intact autophagy induction (Figure S6D-F). Furthermore, LC3-II levels were similar between genotypes, suggesting that autophagosome formation was not impaired (Figure S6G-I). In contrast *CHD2^+/-^* neurons displayed elevated levels of p62 and LAMP1 following autophagy induction, consistent with defective cargo degradation and lysosomal dysfunction (Figure S6G-I). Our results indicate that the reduced autophagosome flux in *CHD2^+/-^*cells primarily arise from impaired lysosomal degradation rather than defective autophagosome biogenesis. Strikingly, autophagosome flux was accelerated in chimpanzee and bonobo cells compared to all lines, with rates reaching approximately 45 autolysosomes per cell per hour (Figure 3K-L), revealing marked interspecies differences in protein degradation capacity. Across species and genotypes, autophagosome flux scaled with CHD2 expression levels, suggesting that CHD2 dosage may contribute to these differences. To directly test this, we ectopically expressed CHD2 in control cortical neurons and observed a positive correlation between CHD2 expression levels and autolysosome abundance (Figure 3M-O), supporting a dosage-dependent relationship between CHD2 and autolysosomal activity. Together, these findings establish CHD2 dosage as a critical regulator of autolysosomal function.

### CHD2 dosage sets the tempo of neuronal maturation via the lysosomal pathway

The bidirectional effects of CHD2 dosage on lysosomal function prompted us to ask whether lysosomal activity contributes directly to the pace of neuronal maturation. To test this possibility, we pharmacologically manipulated the lysosomal pathway in opposite directions. We used SB202190, which induces *autophagy*- and *lysosomal* gene transcription through the translocation of the transcription factor TFEB and TFE3 from the cytosol into the nucleus^38^, and bafilomycin A1 (BafA1), which inhibits lysosomal degradation (Figure 4A). Consistent with its reported mechanism of action^38^, SB202190 increased nuclear TFEB localization in *Ctrl* neurons (Figure S7A). This was accompanied by an increase in autolysosome abundance, indicative of enhanced lysosomal activity. In contrast, BafA1 inhibits the formation of autolysosomes, leading to reduced autophagosome flux (Figure 4B-C). Strikingly, increased autolysosomal activity promoted dendritic growth and complexity, whereas lysosomal inhibition suppressed dendritic development (Figure 4D). These findings indicate that the modulation of lysosomal activity is sufficient to alter neuronal maturation.

**Figure 4.**
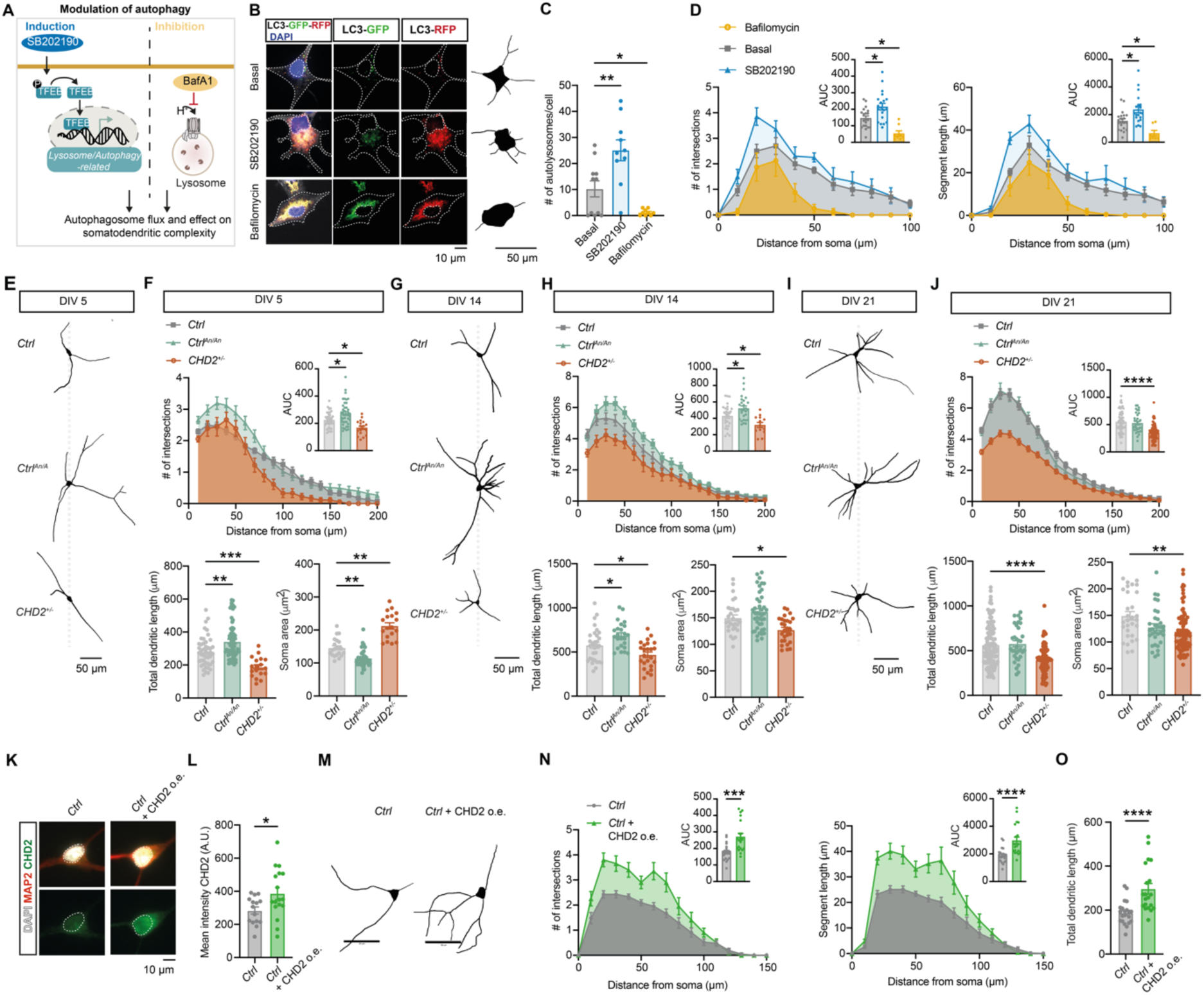
Autolysosomal activity mediates CHD2 dosage-dependent control of neuronal maturation. (A) Schematic overview of pharmacological modulation of the autophagy-lysosome pathway. TFEB activation by SB202190 (10 μM, 4h treatment at the DIV 2) activates transcription of *lysosome*- and *autophagy*-related genes, whereas bafilomycin A1 (BafA1; 200 nM, 15 min. treatment at DIV 2) inhibits lysosomal acidification and autophagic degradation. B) Representative images of tandem LC3-GFP-RFP reporter in DIV 5 cortical neurons treated with SB202190 or BafA1, together with representative neuronal reconstructions from the same conditions. Autolysosomes are identified as GFP-negative/RFP-positive puncta. Scale bars, 10 μm (LC3-GFP-RFP reporter images) and 50 μm (neuronal reconstructions). (C) Quantification of the number of autolysosomes following pharmacological modulation of lysosomal function (n = 11 for basal, n = 10 for SB202190, n = 10 for BafA1). (D) Sholl analysis of dendritic complexity in control cortical neurons following SB202190 or BafA1 treatment. Insets show area under the curve (AUC) (n = 11 for basal, n = 10 for SB202190, n = 10 for BafA1). (E-F) Representative neuronal reconstructions (E) and morphometric quantification (F) of *Ctrl*, *Ctrl^An/An^*, and *CHD2^+/-^*-derived cortical neurons at DIV 5. Quantified parameters include Sholl intersections, total dendritic length, and soma size. Insets show area under the curve (AUC) (n = 49 for *Ctrl*, n = 41 for *Ctrl^An/An^*, n = 18 for *CHD2^+/-^*). (G-H) Representative neuronal reconstructions (G) and morphometric quantification (H) of *Ctrl*, *Ctrl^An/An^*, and *CHD2^+/-^* -derived cortical neurons at DIV 14. Quantified parameters include Sholl intersections, total dendritic length, and soma size. Insets show area under the curve (AUC) (n = 27 for *Ctrl*, n = 30 for *Ctrl^An/An^*, n = 15 for *CHD2^+/-^*). (I-J) Representative neuronal reconstructions (G) and morphometric quantification (H) of *Ctrl*, *Ctrl^An/An^*, and *CHD2^+/-^*-derived cortical neurons at DIV 21. Quantified parameters include Sholl intersections, total dendritic length, and soma size. Insets show area under the curve (AUC) (n = 58 for *Ctrl*, n = 33 for *Ctrl^An/An^*, n = 81 for *CHD2^+/-^*). (K) Representative images of CHD2 immunostaining (green), MAP2 (red) and DAPI (white) in cortical neurons. Scale bar, 10 μm. (L) Quantification of CHD2 expression relative to non-transfected control neurons (n = 18 for *Ctrl* and n = 17 for *Ctrl* +CHD2o.e.) (M) Representative neuronal reconstructions of control and control neurons with ectopic expression of CHD2. Scale bar, 50 μm. (N) Sholl analysis of dendritic complexity in control and CHD2-overexpressing cortical neurons. Insets show area under the curve (AUC) (n = 22 for *Ctrl* and n = 23 for *Ctrl* +CHD2o.e). (O) Quantification of total dendritic length in control and CHD2-overexpressing cortical neurons (n = 22 for *Ctrl* and n = 23 for *Ctrl* +CHD2o.e). Data represent means ±SEM. *p<0.05, **p<0.01, ***p<0.001, ****p<0.0001, one-way ANOVA with post hoc Bonferroni correction, and unpaired two-tailed student t-test.

Previous studies have shown that differences in developmental timing between great apes and modern humans are reflected in the pace of somatodendritic maturation^39^. Given these results, we hypothesized that CHD2-dependent differences in lysosomal activity contribute to developmental timing. We therefore quantified somatodendritic maturation across multiple stages of neuronal differentiation (DIV 5, 14, 21) in *Ctrl^An/An^* and *CHD2^+/-^* neurons relative to their respective controls (Figure 4E-J). Consistent with their enhanced autolysosomal activity, *Ctrl^An/An^* neurons exhibited increased dendritic complexity and neurite length at early developmental stages (DIV 5 and DIV 14) compared to *Ctrl* neurons (Figure 4E-H). Notably, these differences largely disappeared by DIV 21, suggesting a temporal acceleration rather than a sustained increase in dendritic development (Figure 4I-J). Indeed, the dendritic complexity of DIV 14 *Ctrl^An/An^* neurons was comparable to that of DIV 21 *Ctrl* neurons (Figure S7B-C), supporting the notion that elevated CHD2 dosage advances progression through specific stages of neuronal maturation. This relationship was not observed between DIV 5 *Ctrl^An/An^* neurons and DIV 14 *Ctrl* neurons (Figure S7D-E), indicating that CHD2 does not impose a fixed temporal shift, but rather modulates the rate of maturation during defined developmental windows. In contrast, *CHD2^+/-^* neurons displayed reduced dendritic complexity and neurite length across developmental stages, indicative of globally delayed neuronal maturation (Figure 4E-J). Together, these findings reveal a CHD2 dosage-dependent heterochronic shift in neuronal maturation, whereby increased CHD2 levels advances and reduced CHD2 levels delay progression through dendritic developmental states.

To determine whether CHD2 itself is sufficient to promote neuronal maturation, we ectopically expressed CHD2 in control cortical neurons and assessed dendritic development at DIV 5 (Figure 4K-O). CHD2 overexpression resulted in increased dendritic complexity and neurite length, similar to what we observed in *Ctrl^An/An^* neurons (Figure 4M-O, Figure S7F). Together with our observations that elevated CHD2 dosage enhances autolysosomal activity (Figure 3) and that pharmacological activation of the lysosome pathway promotes neuronal maturation (Figure 4A-D), these results support a model in which CHD2 regulates the tempo of cortical neuron development through its effects on lysosomal activity and degradative capacity.

### CHD2 dosage controls developmental tempo in cortical organoids and neuronal networks

The opposing effect of increased and decreased CHD2 dosage on neuronal maturation suggest that CHD2 may act as a regulator of developmental tempo. We therefore asked whether these temporal differences extend beyond individual neurons to influence cortical development at the tissue level. To address this, we performed single-cell RNA sequencing of *Ctrl^An/An^* and *CHD2^+/-^* cortical organoids and compared these to control organoids at Days 25, 50 and 90 of differentiation and reconstructed developmental trajectories across cortical lineages (Figure 5A-D). Immunostainings confirmed that organoids from all genotypes displayed the expected cytoarchitecture and spatial distribution of developmental within the ventricular and subventricular areas (Figure 5A). Leiden clustering was performed to identify cell states and to group them into coherent cell types across the two datasets (Figure S8A-C). We leveraged Leiden-specific differential expression and traced the expression of key cortical markers to annotate cell types (Figure S9A-B). Draw graphs were generated by leveraging PAGA algorithm and compared with diffusion maps to reconstruct differentiation trajectories and lineages (Figure 5B, Figure S9C). Longitudinal analysis of cell-type composition revealed genotype-dependent shifts in lineage progression (Figure 5C). *CHD2*^+/-^ organoids exhibited a consistent depletion of radial glia accompanied by an expansion of intermediate progenitors and mature excitatory neurons expressing markers such as FOXP2 and NEUROD6. These findings are consistent with previous observations in *Chd2*-haploinsufficient mouse models^40^. Our findings are also consistent with the impact of endolysosome inheritance during cell division on radial glia and intermediate progenitors^41,42^. *Ctrl^An/An^* organoids exhibited similar (albeit less pronounced) trends as *CHD2*^+/-^ organoids at Day 25 (Figure 5C). However, at later time points, salient directional changes emerged, with an increase in radial glia in *Ctrl^An/An^*, and, especially at Day 90, a significant increase of (deep-layer) neurons expressing TBR1 and NEUROD6. Given these differences, we quantified the proportion of proliferative cells in G1, G2M, and S phases of the cell cycle compared to *Ctrl* organoids within cyclic progenitors, radial glia, and intermediate progenitors (Figure 5D). On Day 25, G1 phase proportions were similar between *Ctrl^An/An^* and *CHD2*^+/-^ organoids. G2M cells increased by 20% in *Ctrl^An/An^* but decreased by 20% in *CHD2^+/-^* organoids. By Day 50, the proportion of G2M and S phase cells was significantly higher in *Ctrl^An/An^*organoids and considerably lower in *CHD2*^+/-^ ones, indicating that reduced CHD2 dosage accelerates differentiation and depletes the proliferative pool, while increased dosage boosts neurogenesis, with a marked effect on deep-layer neuron production.

**Figure 5.**
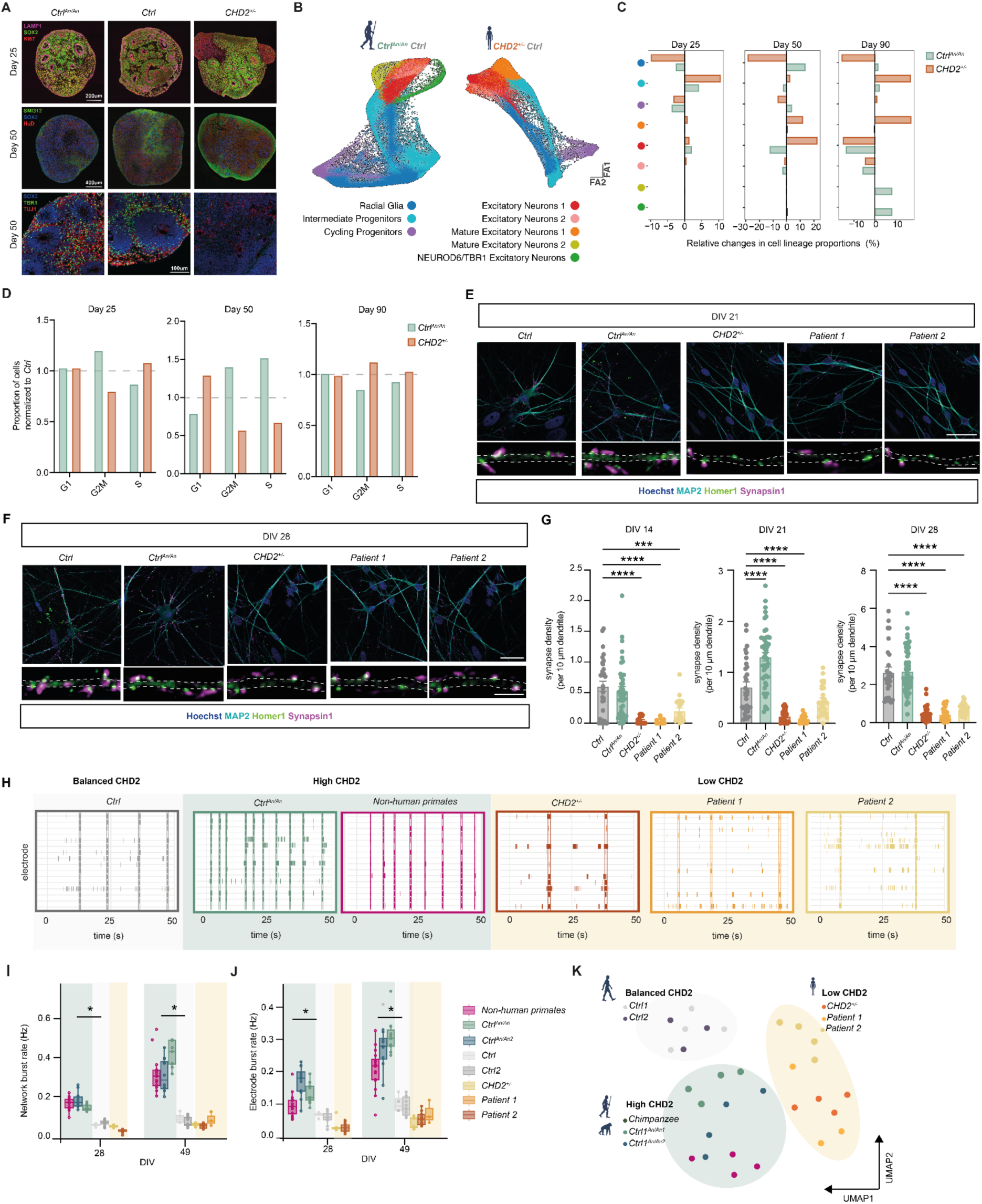
CHD2 dosage regulates the tempo of cortical development and neuronal network maturation. (A) Representative immunocytochemistry staining images of organoids showing relevant markers for lysosomes (LAMP1), proliferation of neuronal progenitors (KI67, SOX2) and cortical development (SMI312, HuD, TBR1 and TUJ1) at Day 25 and Day 50. Scale bar, 200 µm for Day 25, 400 µm for Day 50, 100 µm for zoom in image Day 50. (B) Force-directed graphs of high-quality cells annotated by their lineage in *Ctrl^An/An^* and *CHD2^+/-^*. (C) Butterfly plot showing relative changes in lineage proportions between *CHD2^+/-^* or *Ctrl^An/An^*conditions and the respective *Ctrl*from the same experiment. For each lineage, bars represent the difference in proportion (%). (D) Bar plot of normalized ratios of cell cycle phase proportions (%) normalized to *Ctrl*. Ratios above or below 1 indicate a higher or lower proportion compared to *Ctrl*, respectively. (E) Representative images of immunocytochemistry stained for glutamatergic synapses: Synapsin I as a presynaptic marker and Homer1B/C as a postsynaptic marker at DIV 21. Scale bar, 40 µm for overview image, 4 µm for zoom in image. (F) Representative images of immunocytochemistry stained for glutamatergic synapses: Synapsin I as a presynaptic marker and Homer1B/C as a postsynaptic marker at DIV 28. Scale bar, 40 µm for overview image, 4 µm for zoom in image. (G) Quantification of the density of co-localized Synapsin I/Homer1B/C puncta (number per 10 µm dendrite). Each dot represents the quantification of one cell; (DIV 14 n = 31 for *Ctrl*, n = 41 for *Ctrl^An/An^*, n = 21 for *CHD2^+/-^*, n = 22 for *patient 1*, n = 22 for *patient 2*; DIV 21 n = 30 for *Ctrl*, n = 48 for *Ctrl^An/An^*, n = 25 for *CHD2^+/-^*, n = 18 for *patient 1*, n = 28 for *patient 2*; DIV 28 n = 27 for *Ctrl*, n = 63 for *Ctrl^An/An^*, n = 31 for *CHD2^+/-^*, n = 30 for *patient 1*, n = 30 for *patient 2*). (H) Representative raster plots (50 s) of electrophysiological activity measured by MEA from balanced CHD2 group (*Ctrl*), high CHD2 group (non-human primates [chimpanzee] and *Ctrl^An/An^*) and low CHD2 group *(CHD2^+/-^*, *patient 1*, *patient 2*) -derived neuronal networks at DIV 49. (I-J) Quantification of network (I) and electrode (J) parameters as indicated from DIV 28 and DIV 49. Group with high CHD2 expression (*Ctrl^An/An^* and non-human primates - indicated in green box) and low CHD2 expression (*CHD2^+/-^, patient 1*, and *patient 2* – indicated in yellow box) were compared to the group with balanced CHD2 (*Ctrl1* and *Ctrl2* – indicated in light gray box). Each dot represents the quantification of one well of a MEA plate; (DIV 28 n = 2 for *Ctrl1*, n = 8 for *Ctrl2*, n = 11 for non-human primates [chimpanzee], n = 7 for *Ctrl^An/An^*, n = 10 for *Ctrl^An/An^*^2^, n = 3 for *CHD2^+/-^*, n = 4 for *patient 1*; n = 3 for *patient 2*). (K) UMAP analysis based on 27 MEA parameters across three groups (balanced CHD2 [*Ctrl1* and *Ctrl2*], high CHD2 [*chimpanzee, Ctrl^An/An^*] and low CHD2 [*CHD2^+/-^,* patient 1, patient 2]. Data represent means ±SEM. *p<0.05, **p<0.01, ***p<0.001, ****p<0.0001, one-way ANOVA with post hoc Bonferroni correction.

Because developmental timing ultimately influences the establishment of neuronal circuits, we next investigated the impact of CHD2 dosage on synapse formation and network maturation. Ctrl*^An/An^* neurons exhibited an increase in functional excitatory synapses at DIV 21, as indicated by an increased number of co-localizing synapsin1 and homer1B/C puncta (Figure 5E,G). Consistent with the transient dendritic phenotype, these differences largely disappeared by DIV 28 (Figure 5F-G). In contrast, *CHD2* haploinsufficient neurons (*CHD2^+/-^,* patient 1 and patient 2) exhibited a marked reduction in excitatory synapse density that persisted throughout maturation (Figure 5E-G).

To determine whether these cellular phenotypes translate into altered network function, we recorded spontaneous activity from *Ctrl^An/An^*, *CHD2* haploinsufficient (*CHD2^+/-^,* patient 1 and patient 2) and chimpanzee neuronal cultures using microelectrode arrays (MEAs) (Figure 5H-K). By DIV 28, *Ctrl* cultures developed synchronized network bursts characteristics of functionally connected neuronal networks. In line with their accelerated maturation, chimpanzee and *Ctrl^An/An^* cultures displayed elevated firing and network burst rates compared to *Ctrl*, whereas *CHD2* haploinsufficient cultures showed reduced network activity (Figure 5H-J). Consistent with these opposing functional phenotypes, UMAP analysis of all independent MEA network parameters segregated high-CHD2 and low-CHD2 networks into distinct clusters, with chimpanzee-derived cultures grouping closely with *Ctrl^An/An^*-derived networks (Figure 5K). Together, these findings demonstrate that CHD2 dosage influences developmental tempo across multiple functional scales.

### Lysosomal modulation restores neuronal maturation in *CHD2^+/-^* neurons

To establish whether lysosomal dysfunction contributes causally to impaired neuronal maturation in *CHD2^+/-^* neurons, we designed complementary genetic and pharmacological rescue strategies targeting the lysosomal pathway (Figure 6A). We first sought to identify molecular components of the lysosomal machinery that are altered upon *CHD2* haploinsufficiency and could therefore serve as targets for lysosomal restoration. Given that lysosomal acidification depends on proper assembly and function of v-ATPase proton pumps, we examined whether components of this machinery were also affected in *CHD2^+/-^* neurons. In *Ctrl* neurons, protein levels of NCOA7 along with ATP6V1A and ATP6V1B2 protein levels, two essential components of the v-ATPase complex, progressively increased as neurons matured, consistent with the developmental acquisition of lysosomal function (Figure 6B, Figure S10A). In contrast, *CHD2^+/-^* neurons exhibited a markedly attenuated increase in all three proteins, which remained at lower levels throughout differentiation (Figure 6B). These findings suggest that *CHD2* haploinsufficiency delays acquisition of the lysosomal machinery required for acidification.

**Figure 6.**
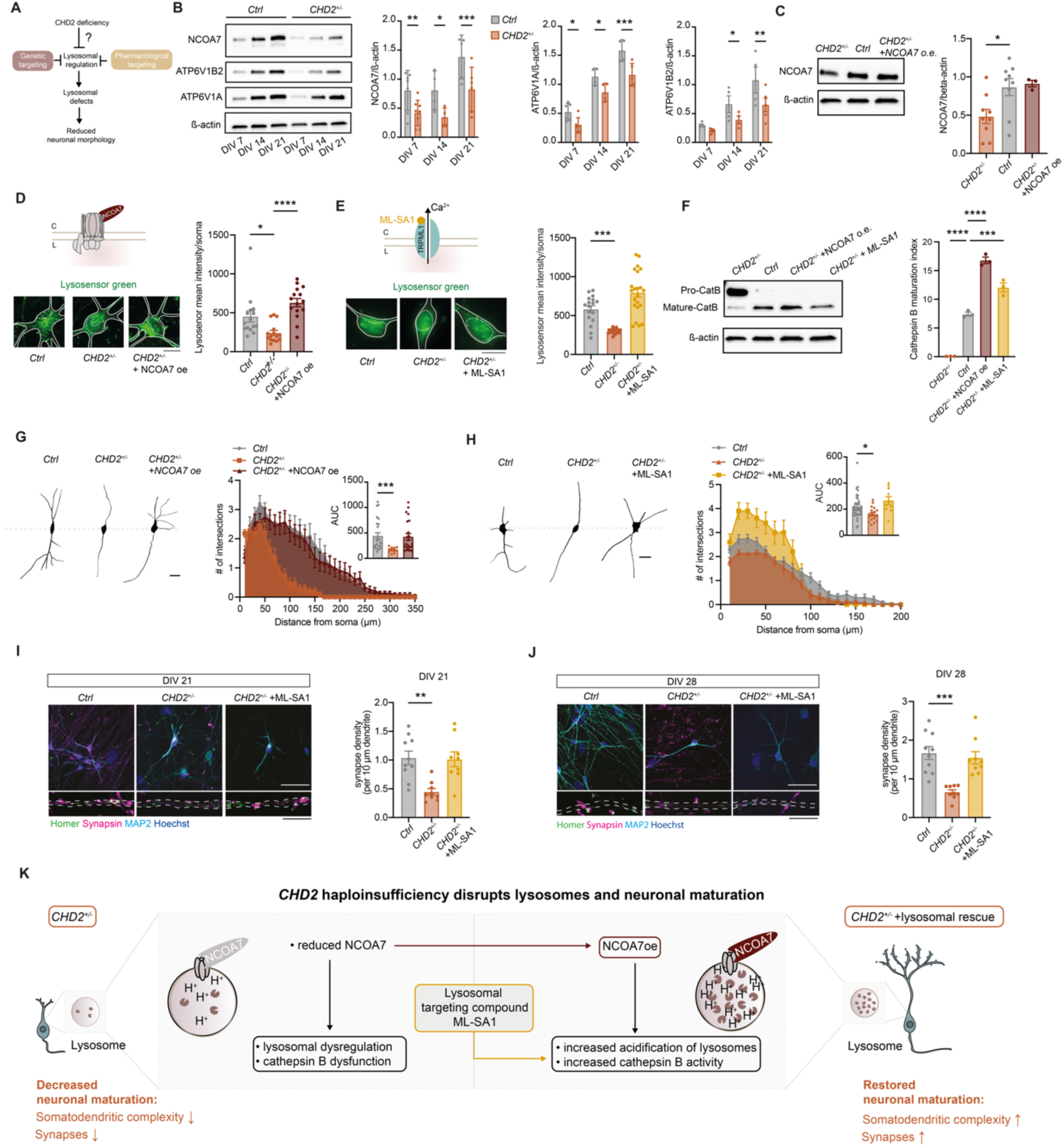
NCOA7 and lysosome-targeting compounds restore lysosomal function and neuronal maturation in *CHD2*-haploinsufficient neurons. (A) Schematic overview of the rescue strategy. Candidate lysosomal regulators identified downstream of CHD2 deficiency were targeted genetically or pharmacologically to restore lysosomal function and neuronal maturation. (B) Western blot and quantification of NCOA7, ATP6V1B2, and ATP6V1A expression in control and *CHD2^+/-^* -derived cortical neurons across differentiation (DIV 7, DIV 14, DIV 21) (n = 5 for *Ctrl*, n = 5 for *CHD2^+/-^*). (C) Western blot and quantification confirming NCOA7 overexpression (o.e.) in *CHD2^+/-^* cortical neurons (n = 9 for *CHD2^+/-^*, n = 9 for *Ctrl*, n = 3 for *CHD2^+/-^* +NCOA7o.e.). (D) Representative LysoSensor green images and quantification of LysoSensor green intensity in *Ctrl*, *CHD2^+/-^*, and *CHD2^+/-^* + NCOA7o.e. neurons. Scale bar, 10 µm (n = 16 for *Ctrl*, n = 15 for *CHD2^+/-^*, n = 15 for *CHD2^+/-^* +NCOA7o.e.). (E) Representative LysoSensor green images and quantification of LysoSensor green intensity following treatment ML-SA1 (25 μM for 15 min). Scale bar, 10 µm (n = 17 for *Ctrl*, n = 15 for *CHD2^+/-^*, n = 22 for *CHD2^+/-^* +NCOA7o.e.). (F) Western blot and quantification of cathepsin B maturation index following NCOA7 overexpression or ML-SA1 treatment (n = 3 for *Ctrl*, n = 3 for *CHD2^+/-^*, n = 3 for *CHD2^+/-^* +NCOA7o.e., n = 3 for *CHD2^+/-^* +ML-SA1). (G) Representative neuronal reconstructions and Sholl analysis of *Ctrl*, *CHD2^+/-^*, and *CHD2^+/-^* +NCOA7o.e. treated neurons at DIV 7. Scale bar, 50 µm. Insets show area under the curve (AUC) (n = 30 for *Ctrl*, n = 25 for *CHD2^+/-^*, and n = 25 for *CHD2^+/-^* +NCOA7o.e). (H) Representative neuronal reconstructions and Sholl analysis of *Ctrl*, *CHD2^+/-^*, and *CHD2^+/-^* +ML-SA1 neurons at DIV 7. *CHD2^+/-^*neurons were treated with 25 μM every other day). Scale bar, 50 µm. Insets show area under the curve (AUC) (n = 30 for *Ctrl*, n = 30 for *CHD2^+/-^*, and n = 30 for *CHD2^+/-^* +ML-SA1). (I) (left) Representative images of immunocytochemistry stained for glutamatergic synapses: Synapsin I as a presynaptic marker and Homer1B/C as a postsynaptic marker at DIV 21. Scale bar, 40 µm for overview image, 4 µm for zoom in image. (right) Quantification of the density of co-localized Synapsin I/Homer1B/C puncta, from 3 independent replicates (n = 10 for *Ctrl*, n = 10 for *CHD2^+/-^*, n = 10 for *CHD2^+/-^* +ML-SA). (J) (left) Representative images of immunocytochemistry stained for glutamatergic synapses: Synapsin I as a presynaptic marker and Homer1B/C as a postsynaptic marker at DIV 28. Scale bar, 40 µm for overview image, 4 µm for zoom in image. (right) Quantification of the density of co-localized Synapsin I/Homer1B/C puncta, from 3 independent replicates (n = 10 for *Ctrl*, n = 10 for *CHD2^+/-^*, n = 10 for *CHD2^+/-^* +ML-SA). (J) Working model illustrating how *CHD2* haploinsufficiency reduces expression of lysosomal regulators, resulting in impaired lysosomal acidification, defective cathepsin B maturation, and delayed neuronal maturation. Genetic (NCOA7 overexpression) or pharmacological restoration of lysosomal function (ML-SA1) rescues degradative capacity and neuronal maturation. Data represent means ±SEM. *p<0.05, **p<0.01, ***p<0.001, ****p<0.0001, one-way ANOVA with post hoc Bonferroni correction.

Given the reduced expression of NCOA7 in *CHD2^+/-^* neurons and its established role in promoting lysosomal acidification functioning as a scaffold for the v-ATPase assembly^43^, we next tested whether restoring NCOA7 protein levels was sufficient to rescue the lysosomal defects in *CHD2*-haploinsufficient neurons. Overexpression of NCOA7 in *CHD2^+/-^* neurons restored lysosomal pH (Figure 6C-D). We further assessed whether lysosomal acidification could also be restored pharmacologically. To this end, *CHD2^+/-^* neurons were treated with ML-SA1, a small-molecule agonist of the lysosomal calcium channel TRPML1 that has been shown to enhance lysosomal function^44,45^ (Figure 6E). Similar to NCOA7 overexpression, ML-SA1 treatment significantly increased LysoSensor fluorescence in *CHD2^+/-^*neurons, restoring lysosomal acidification to levels comparable to those observed in *Ctrl* neurons (Figure 6E). The convergence of NCOA7 overexpression and ML-SA1 treatment on lysosomal acidification provided two independent approaches to test whether restoration of lysosomal pH is sufficient to recover downstream proteolytic function. Because our previous analyses identified cathepsins as downstream targets of CHD2 dosage, and specifically the expression and activity of cathepsin B altered by CHD2 expression (Figure 3C), we tested whether restoration of lysosomal acidification could also rescue cathepsin maturation. Both NCOA7 overexpression and ML-SA1 treatment significantly increased cathepsin B maturation in *CHD2^+/-^* neurons (Figure 6F).

We next asked whether restoration of lysosomal acidification and cathepsin B maturation translates into improved neuronal maturation. To address this, we reconstructed individual neurons following NCOA7 overexpression or treatment with ML-SA1 and quantified dendritic complexity using Sholl analysis (Figure 6G-H). NCOA7 overexpression significantly increased dendritic arborization and restored Sholl profiles toward *Ctrl* levels (Figure 6G). Similarly, treatment with the lysosome-modulating compound ML-SA1 enhanced dendritic complexity in *CHD2^+/-^* neurons (Figure 6H). To determine whether this rescue extended beyond dendritic morphology to synaptic maturation, we next quantified synapse density following restoration of lysosomal function by ML-SA1 at DIV 21 and DIV 28. ML-SA1 treatment lead to an increased number of co-localizing synapsin1 and homer1B/C puncta in *CHD2^+/-^* neurons, restoring it to levels comparable to *Ctrl* neurons (Figure 6I-J). Together, these findings indicate that pharmacological enhancement of lysosomal function is sufficient to improve neuronal maturation.

Based on these findings, we conclude that impaired lysosomal acidification and cathepsin B maturation are mechanistically linked to the neuronal maturation defects caused by *CHD2* haploinsufficiency. Restoring lysosomal function, either genetically through NCOA7 overexpression or pharmacologically through a lysosome targeting compound, rescues dendritic and synaptic development in *CHD2^+/-^* neurons (Figure 6K). These results identify lysosomal dysfunction as a therapeutically tractable mechanism downstream of *CHD2* haploinsufficiency.

## DISCUSSION

By integrating the study of neurodevelopmental disorders with the analysis of evolutionary variants, we here provide evidence that CHD2 dosage is a critical arbiter of human cortical maturation. We demonstrate that a *sapiens*-specific enhancer variant reduces *CHD2* expression, whereas reversion to the ancestral allele promotes earlier neuronal maturation. Across isogenic, patient-derived, and non-human primate models, we show that CHD2 dosage bidirectionally regulates lysosomal acidification, neurite outgrowth, and network activity. These findings establish lysosomal function as a fundamental determinant of developmental tempo, reinforcing the emerging role of lysosomal dynamics in instructing neural stem cell fate and neuronal maturation^36,41^. From an evolutionary perspective, our results link increased lysosomal function to accelerated cortical maturation, identifying the autophagy-lysosome pathway as a key cellular trait shaping species-specific developmental programs.

Although lysosomes have traditionally been viewed as passive degradative organelles, our study establishes the autophagy-lysosome pathway as an active regulator of cortical development and neuronal maturation. Recent evidence has highlighted how the asymmetric inheritance of lysosomes during cell division influences the fate and proliferative capacity of radial glia and intermediate progenitors^41^, while autophagy supports the extensive organelle remodeling required for the transition from progenitors to post-mitotic neurons^36,41^. Once specified, the spatial organization and positioning of lysosomes within dendrites further facilitate the local protein turnover necessary for dendritic development and synaptic remodeling^46^. Our findings extend this by placing CHD2 upstream of these processes. Altered CHD2 dosage reciprocally modulates lysosomal acidification and the maturation of cathepsin B, a protease required for neurite outgrowth^47^. Furthermore, direct manipulation of the lysosomal pathway was sufficient to bidirectionally tune dendritic complexity, demonstrating that lysosomal degradative capacity actively instructs structural maturation. Critically, these lysosomal defects are not merely downstream correlates of *CHD2* loss but causal drivers of the phenotype, as we were able to demonstrate that restoring lysosomal acidification, either via NCOA7 overexpression or pharmacological agents, rescued maturation in *CHD2*-haploinsufficient neurons. These data thereby also identify lysosomal acidification as a candidate therapeutic entry point for *CHD2*-associated neurodevelopmental disorders.

Our findings exemplify how a single regulatory variant can exert a significant causative role in setting divergent developmental courses in closely related species *in vitro*. But a few points bear emphasis in the context of our study. First, although the regulatory variant we examined yields reliable effects mirroring those found in *CHD2*^+/-^ lines, *CHD2* haploinsufficiency produces a pronounced disruption of lysosomal function and neuronal maturation, whereas the ancestral enhancer variant in *CHD2* induces comparatively modest but highly reproducible changes. This nicely illustrates both the evolutionary potential of regulatory changes, but also the more context-sensitive and developmental-stage-dependent nature of such changes, as opposed to protein-coding changes. Notably, the functional changes caused by the ancestral enhancer variant were also often transient; the somatodendritic development and synapse formation phenotypes in ancestralized neurons eventually converged with controls, supporting a heterochronic interpretation where CHD2 dosage modulates the rate of progression through maturational states rather than final structural complexity. By contrast, *CHD2* haploinsufficiency resulted in persistent developmental delays. This transient nature of the phenotypes observed here has broader implications: early perturbations in lysosomal activity may initiate developmental cascades that result in substantial downstream differences, even after the initiating phenotype has cleared. This principle of “developmental priming” may be especially relevant for dosage-sensitive chromatin remodelers, where subtle early shifts in gene expression are propagated across trajectories. In this respect, our findings parallel work on other genes studied for their role in neurodevelopment such as like *CHD8*^48–51^, highlighting altered developmental tempo rather than static cellular states as a central disease-relevant phenotype.

Naturally, as no variant or gene is an island, our findings are to be interpreted in a broader context, with the modern variant we dissected and the nature of its target gene (a chromatin remodeler), constituting an experimentally tractable component of a complex evolutionary network. For this reason, our results lead us to anticipate additional changes in genes implicated in lysosomal dynamics and autophagy-lysosome pathways to show important differences between us and our extinct relatives. Genes like *RB1CC1* ^52^, *SETD1A*^53^, and *TFEB*^54^, already highlighted in the paleogenomic literature^29^, are strong candidates for further study. Our results also lead us to predict synergistic changes in domains like the cell cycle and neurogenesis, already a focus of the ancient DNA literature^19^. More generally, our results point to the value – and feasibility – of prioritizing, from the vast amount of *sapiens* nearly fixed regulatory variants and the much smaller core of protein-coding changes, specific combinations to be probed for their functional synergy. We recently spearheaded such perturbations in neural organoids of 15 genes in all pairs of possible combinations to reconstruct reciprocal regulatory hierarchies^55^. A case in point is the nearly-fixed missense mutation that was acquired in *SLITRK1* during the evolution of *Homo sapiens* and that leads to neurite related changes *in vitro*^56^. This gene is among the DEGs in our Day 50 organoids, and its interaction with *CHD2* and other deep-layer cortical markers that are also DEGs in our study, like *FOXP2* and *NFIB*. These too also harbor *sapiens*-derived changes in enhancer regions active during early cortical development^29^, could thus provide an ideal entry point to explore this research question further.

Finally, our results must be placed in a broader evolutionary context, beyond the *Homo* lineage. Changes in neuronal maturation distinguishing *Homo* from *Pan* have featured prominently in the literature^57,58^, and the transient effects we obtained may be seen as part of a “neoteny” continuum. Consistent with this interpretation, the effects of the ancestral enhancer were generally smaller than the differences observed between *sapiens* and non-human primate neurons, indicating that additional evolutionary modifications likely contribute to species-specific regulation of CHD2 and the lysosomal pathway. Crucially, this evolutionary stretching of developmental and maturation window appears to have been selected at the cost of developmental robustness. In this light, our findings resonate with emerging evidence that the evolutionary changes in the expression of dosage-sensitive ASD genes may have increased susceptibility to neurodevelopmental disorders in modern humans. A recent comparative single-cell transcriptomic analysis proposed that human cortical neurons, particularly the abundant layer 2/3 intratelencephalic neurons, underwent coordinated downregulation of a broad set of ASD-associated genes during human evolution, potentially as a consequence of positive selection^59^. They proposed that this shift moved the expression of dosage-sensitive ASD genes closer to a critical functional threshold, thereby increasing vulnerability to subsequent genetic or environmental perturbations. Our identification of a modern human-specific enhancer variant that reduces CHD2 expression provides a mechanistic example supporting this broader framework. Because *CHD2* is a dosage-sensitive ASD-linked gene, and our data demonstrate that even modest reductions in its expression are sufficient to perturb lysosomal function, reprogram developmental trajectories, and impair neuronal maturation. Together, these findings support a model in which evolutionary fine-tuning of dosage-sensitive neurodevelopmental regulators contributed to species-specific developmental programs while simultaneously reducing the buffering capacity of human neuronal systems, increasing the likelihood that subsequent perturbations push gene expression below a pathogenic threshold.

Taken together, our findings support a model in which CHD2-dependent regulation of lysosomal functions controls the tempo of neuronal maturation. Our study establishes lysosomes as active regulators of developmental timing rather than passive degradative organelles. Importantly, decoding this evolutionary trajectory bears significant translational implications, uncovering lysosomal homeostasis as a mechanistically grounded and tractable therapeutic vulnerability for neurodevelopmental disorders caused by CHD2 haploinsufficiency.

### Material and Methods

### Generation of human induced pluripotent stem cells

All experiments involving material from patients were carried out after informed consent was obtained and with approval from the medical ethical committee of Radboud University Medical Center, Nijmegen (2018 – 4525). Peripheral mononuclear blood cells (PBMCs) were isolated from patients via blood samples taken during routine diagnostic testing. CHD2 patient 1 was a male patient who had a heterozygous *de novo* mutation in exon 17 (c.2077_2078dup p.Glu694*). CHD2 patient 2 was a female patient with a missense mutation in exon 9 (c.948C>A p.Tyr316*). The clinical history of all patients is described in the Supplementary table 1. All lines were reprogrammed using episomal vectors with the Yamanaka transcription factors Oct4, c-Myc, Sox2 and Kfl4 and had normal karyotypes.

### CRISPR/Cas9 gene editing of *CHD2*

CRISPR/Cas9 technology was used to create a heterozygous indel variant in exon 4 of *CHD2* in two independent human iPS cell lines derived from a healthy male and female donor, which were obtained from the Corriel Institute (USCFi001-A, GM25256, indicated as *Ctrl1* in this study) and Center for iPS cell Research and Application (CIRAAi007-A, indicated as *Ctrl2* in this study). In brief, a short guide RNA (sgRNA) was designed (ATCCTGATGTTTATGGGGTC) and cloned into pSpCas9(BB)-2A-Puro (PX459) V2.0 (Addgene, #62988) according to previous studies. 8×10^5^ single hiPSCs were nucleofected with 5 µg of the generated pSpCas9-sgRNA plasmid using the P3 Primary Cell 4D-Nucleofector Kit (Lonza, #V4XP-3024) in combination with the 4D Nucleofector Unit X (Lonza, #AAF-1002X). After nucleofection, cells were resuspended in E8 Flex supplemented with RevitaCell (Thermo Fisher Scientific, #A2644501) and seeded on biolaminin 521 (Biolamina, #LN521) pre-coated wells. 24 hours after nucleofection, 0.5 µg/ml puromycin was added for 24h. Puromycin resistant colonies were manually picked and send for Sanger Sequence to ensure heterozygous editing of exon 4. Genomic stability was assessed by detection of recurrent abnormalities using the iCS-digital^TM^ PSC test, provided as a service by Stem Genomics (https://www.stemgenomics.com/) (Figure S2E, Figure S5C). We observed a loss of X chromosome in *CHD2^+/-^* generated in *Ctrl2*, which was also confirmed by karyotyping (Figure S5D). Whole genome sequencing and off-target analysis was performed to confirm genetic integrity after genomic editing (Figure S5E).

### CRISPR/Cas9 gene editing in enhancer region of *CHD2*

Isogenic clones harboring the ancestral allele in the enhancer region were generated from the GM25256 cell line. Briefly, a sgRNA (CACCTGCACTCAGAGTCAGCTTGT) was cloned into BbsI-HF-digested pSpCas9(BB)-2A-Puro (PX459) V2.0 (Addgene, #62988). A single cell suspension of 8×10^5^ human iPS cells was nucleofected with 5 µg of the pSpCas9-sgRNA plasmid and 200 pmol of single-stranded oligodeoxynucleotides (ssODN; sequence: CTTTTTCCCTTGCTGTACCCGGGACACATTGATTTACATTTTAGATCTGCATCTGTGTCAAGATTAGACCAGCTATTCCAGTCAAAGAGTCAGCTTGCATTCAGAGTCAGCTTGTTGGCCCACCCTTTCCACAGACATAATAATGAATCAGATAGTGGGAAGCCAGGAGAAGGTATTGTCAAATGCTGGGGGTTAA ATCC) using the P3 Primary Cell 4D-Nucleofector Kit (Lonza, #V4XP-3024) with the 4D Nucleofector Unit X (Lonza, #AAF-1002X). Cells were then resuspended in mTeSR1 (Stemcell Technologies, # 85850) supplemented with 10 µM Y-27632 dihydrochloride (R&D Systems, #1254) and plated on Matrigel (Corning, #354277) pre-coated wells. 24 hours after nucleofection, 0.25 µg/ml puromycin was added for 24h. Puromycin resistant colonies were manually picked and sent for Sanger sequencing to ensure homozygous editing. Genomic stability was assessed through whole genome sequencing (Figure S5E). In the female *Ctrl2* and *Ctrl2^An/An^* lines, we observed a loss of X chromosome (Figure S2E). To check for off-target activity, we leveraged CasOFFinder (10.1093/bioinformatics/btu048) to predict off-targets with up to 3 mismatches (Figure S2C-D) with the sgRNA sequences. We then designed primer for each off-target and performed PCR followed by Sanger sequencing to exclude off-target activity.

### Human induced pluripotent stem cells

All *CHD2*-haploinsufficient iPS cell lines were cultured on biolaminin 521 (Biolamina, #LN521), and *Ctrl^An/An^* lines under feeder-free conditions either on Matrigel (Corning, #354277) coated plates or 0,125mg/cm^2^ of iMatrix-511 (ReproCell, #NP892-012). Chimpanzee derived iPS cells (male) and bonobo derived iPS cells (female) were obtained from MPI EVA from Svante Pääbo lab. Cells were cultured in essential 8 (E8) flex medium (Gibco, #A2858401) supplemented with primocin (0.1 µg/mL, Invivogen, #ant-pm-2) or mTeSR1 (Stemcell Technologies, #85850) medium supplemented with 100 U/mL penicillin and 100 ug/mL streptomycin (Euroclone, #ECB3001D). Cells were passaged before reaching 70% confluency using an enzyme-free reagent (ReLeSR, Stemcell, #05872) or a 0.5 mM EDTA solution and not kept for more than 10 passages. All hiPSCs were regularly tested for mycoplasma contamination using MycoAlert PLUS (Lonza, #LT07-703) and their identity was confirmed by short tandem repeat profiling. To make *Neurogenin 2* (*Ngn2*) doxycycline-inducible excitatory neurons, cells were infected with lentiviral *Ngn2* and *TRE3G* constructs using the transfer vector pLV[TetOn]-Puro-TRE3G>mNeurog2 (Addgene plasmid # 198754) and pLV-EF1A>Tet3G:IRES:Neo (Addgene plasmid # 198756), respectively. Selection was performed by G418 (50 µg/mL, Sigma-Aldrich, #G8168) for *Ngn2*, and by puromycin (0.5 µg/mL) for *TRE3G*. hiPSCs were kept at 37°C/5%/CO_2_. The use of iPSC lines was approved by the ethical committee of the University of Milan, and RadboudUMC Nijmegen.

### *Ngn2* neuronal differentiation

Human iPS cells were differentiated into cortical neurons as previously described^60–62^. Briefly, iPS cells were directly differentiated into excitatory cortical layer 2/3 neurons by overexpressing *Ngn2* upon doxycycline treatment (1 mg/mL). Neuronal maturation was supported by E18 rodent astrocytes, which were added to the culture in a 1:1 ratio two days after iPS cells plating. Cell plating was adjusted to ensure equal cell densities. Well of neuronal cultures that had an unequal density or showed clustering were excluded.

### Cortical brain organoids

Cortical brain organoids were generated as previously described^63^. Briefly, 2×10^4^ iPS cells were plated in 96 V-bottom ultra-low attachment plates (S-Bio, #MS-9096VZ) in 100 µl of stem cell medium supplanted with 5 µm Y-27632 dihydrochloride (R&D Systems, #1254) and centrifuged for 3 minutes at 150 rcf to promote spheroids aggregation. Starting from day 0, medium was replaced daily with neural induction medium containing 80% DMEM/F12 medium (Gibco, #11330057), 20% Knockout serum (Gibco, #10828028), non-essential amino acids 1:100 (Euroclone, #ECB3054D), 0.1 mM cell culture grade 2-mercaptoethanol solution (ThermoFisher Scientific, #31350010), GlutaMax 1:100 (Gibco, #35050061), penicillin 100 U/mL and streptomycin 100 µg/mL (Euroclone, #ECB3001D), 5 µM Dorsomprhin (RayBiotech, #331-21079-2) and 10 µM TGFβ-inhibitor SB431542 (MedChem Express, #HY-10431). On day 6 medium was changed to complete Neurobasal medium (Gibco, #12348017) with B27 supplement without vitamin A 1:50 (Gibco, #12587001), GlutaMax 1:100 (Gibco, #35050061), penicillin 100 U/mL and streptomycin 100 μg/mL (Euroclone, #ECB3001D), 0.1 mM cell culture grade 2-mercaptoethanol solution (ThermoFisher Scientific, #31350010) supplemented with 20 ng/mL FGF2 (Peprotech, #100-18B) and 20 ng/mL EGF (Peprotech, #AF-100-15). On day 12, organoids were transferred to 90mm ultra-low attachment dishes (S-Bio, #MS-90900Z) on a standard orbital shaker (VWR Standard Orbital Shaker, Model 1000) at 55 rpm. From day 12 onwards, medium was changed every other day. On day 25, FGF and EGF were replaced with 20 ng/mL of BDNF (PeproTech, #450-02) and 20 ng/mL neurotrophin-3 (Peprotech, #450-03). From day 42 onwards, complete Neurobasal medium without growth factors was used and changed 3 times a week.

### Western blot

For western blot assays, a total of 350,000 cells were seeded on a six-well plate. Proteins were extracted in cold RIPA lysis buffer consisting of 50 mM Tris HCl (pH 7.5), 150 mM NaCl, 1% NP-40, 0.10% SDS, 0.50% sodium deoxycholate, 1 mM EDTA, diluted in H_2_O, or by NE-PER^TM^ Nuclear and Cytoplasmic Extraction Reagents (Thermo Fisher, #78835). Pierce BCA protein assay kit (ThermoFisher, #23225) was used to determine protein concentration. Protein lysates were loaded on Mini-PROTEAN TGX stain-free gels (Bio-Rad, #4568084) before transfer to 0.2 µm PVDF membrane (Bio-Rad, #1704156), or nitrocellulose (Bio-Rad, #1704158). The membrane was blocked with 5% blotto non-fat dry milk (ChemCruz, #sc-2325) or 5% bovine albumin serum (Sigma, #A7906-100G) (in 1X TBST 0.1%) for 1h at room temperature. Primary antibodies were incubated on the membrane for overnight at 4°C. The next day, membranes were washed three times in TBS-T0.1% followed by incubation with corresponding secondary antibodies for 1h at room temperature. Membranes were washed five times in TBS-T 0.1%, before ECL detection with the SuperSignal^Tm^ West Femto kit (Thermo Scientific, #34095). The membranes were imaged using the Bio-Rad Gel Imaging system (ChemiDoc^TM^ Touch Imaging System Bio-Rad). The following primary antibodies were used: rabbit anti-CHD2 (1:1000, Cell Signaling Technology, #4170S), mouse anti-β-actin (1:5000; Invitrogen, #MA1-140), mTOR (1:1000, Cell Signaling, #2972), p-mTOR (1:1000, Cell Signaling, #2971), ULK1 (1:1000, Cell Signaling, #8054), p-ULK1 (1:1000, Cell Signaling, #14202), LC3 (1:200, NanoTools, 0231-100/LC3-5F10), P62 (1:200; Sigma, p0067), g-tubulin (1:1000, BioLegend, #MMS-435P), NCOA7 (1:1000, Proteintech, #23092-1-AP), ATP6V1A, (1:1000, Proteintech, #17115-1-AP), ATP6V1B2 (1:1000, Proteintech, #15097-1-AP) and cathepsin B (1:1000, Invitrogen, #PA587537). Secondary antibodies, conjugated to horseradish peroxidase (HRP), were diluted in 5% milk and added for 1h at room temperature. The following secondary antibodies were used: goat anti-mouse (1:50 000, Jackson ImmunoResearch Laboratories, 115-035-062) and goat anti-rabbit (1:50 000, Invitrogen, #G21234).

### Quantitative polymerase chain reaction

RNA samples were isolated from iPS cells using the Nucleospin RNA isolation kit (740955, Machery Nagel) according to the manufactures’ instructions. Synthesis of cDNA was performed by iScript cDNA synthesis kit (1708890, Bio-Rad). PCR reactions were done by GoTaq master mix 2x with SYBR Green (A6002, Promega) according to the manufacturer’s protocol. PCR reactions were performed in a 96-well 0.2 ml format (Thermo Fisher Scientific) and ran on the QuantStudio^TM^ Real-Time PCR systems (Thermo Fisher Scientific). All samples were analyzed in duplicate in the same run. Reverse transcriptase-negative conditions and no template-controls were used as a negative control. We used the arithmetic mean of the C_t_ values of the technical replicates to calculate the relative mRNA expression levels of *CHD2*. The mRNA expression level was calculated using the 2^−ΔCt^ method with standardization to PPIA (Peptidylprolyl Isomerase A), GAPDH (glyceraldehyde-3-phosphate dehydrogenase) and TBP (TATA-Box Binding protein). Primers used to quantify mRNA expression of *CHD2* are Fw (5’-3’) CTGTGACAGGTGGGGAAGAG and Rv (5’-3’) TTCCAAGCCATCATCTTTCA. Primers used for housekeeping mRNAs are: PPIA Fw (5’-3’) CATGTTTTCCTTGTTCCCTCC and Rv (5’-3’) CAACACTCTTAACTCAAACGAGGA; GAPDH Fw (5’-3’) GTGGACCTGACCTGCCGTCT and Rv (5’-3’) GGAGGAGTGGGTGTCGCTGT; TBP Fw (5’-3’) GAGCCAAGAGTGAAGAACAGTC and Rv (5’-3’) GCTCCCCACCATATTCTGAATCT.

### Immunocytochemistry

Cultured cells were fixed with 4% paraformaldehyde supplemented with 4% sucrose for 15 min at RT, washed three times with 1XPBS, and permeabilized with 0.2% triton diluted in 1XPBS for 10 min at RT. Nonspecific binding sites were blocked with blocking buffer (5% normal goat serum diluted in 1XPBS) for 1h at room temperature. Primary antibodies were diluted in blocking buffer and incubated overnight at 4°C. Secondary antibodies conjugated to either Alexa 488, Alexa 568 or Alexa 647 (Invitrogen) were incubated for 1h at 1:1000 or 1:300 dilution in blocking buffer for neurons or organoids, respectively. Cell nuclei were stained with Fluoromount-G Mounting medium (with DAPI) at 1:50 000 dilution in 1XPBS for 10 minutes at RT. Organoids collected for imaging were washed twice in PBS and fixed in 4% PFA overnight hours at 4°C and then dehydrated for 24-48 hours in 30% sucrose in PBS 1X. For cryopreservation, organoids are transferred to plastic molds filled with a 1:1 solution of optimal cutting temperature compound (OCT) and 30% sucrose and snap-frozen by immersion in cold isopentane. 15 µm-thick slices were obtained through cryostat sectioning and placed on superfrost glass slices. Antigen retrieval was performed by heating the slices at 70 °C for 40 minutes in Dako Target Retrieval Solution 10X, Citrate Ph6 (Agilent Technologies, S236984-2) diluted 1:10 in PBS. Permeabilization was performed for 30 minutes at room temperature in 0.5% Triton-X in PBS 1x followed by incubation in blocking solution (0.3% Triton-X + 5% Normal Donkey Serum in PBS 1x) at room temperature for 1 hour. The following primary antibodies and dilutions were used: rabbit anti-CHD2 1:1000 (4170S, Cell Signaling Technology); guinea pig anti-MAP2 647 1:1000 or 1:5000 in neurons or organoids, respectively (188004, Synaptic Systems); rabbit anti-Synapsin I 1:500 (AB1543P, Sigma-Aldrich); Homer1b/c 1:200 (160111, Synaptic Systems); mouse anti-LAMP1 1:200 (15665S, Cell Signaling); goat anti-SOX2 1:300 (AF2018, R&D Systems); rabbit anti-TFEB 1:1000(13372-1-AP, Proteintech) rabbit anti-TBR1 1:200 (ab183032, Abcam), mouse anti-SMI312 1:300 (837904, Biolegend), rabbit anti-HuD 1:200 (ab184267, Abcam), mouse anti-KI67 1:100 (9449S, Cell Signaling), mouse anti-TUJ1 (Tubulin beta 3) 1:300 (801202, Biolegend).

### Autophagosome flux assay

iPSCs were transfected with a lentiviral vector expressing tandem mCherry/GFP-LC3 for 7h, and then washed with 1XPBS. Two days after transfection, medium was refreshed 3h prior to rapamycin treatment. Cells were treated with 200 nM rapamycin (sc-3504A, Santa Cruz Biotechnology) for 10 min. at 37°C/CO_2_ and washed with 1XPBS. After 3h, 5h, and 7h of autophagy induction, the cells were fixed with 4% PFA/sucrose diluted in 1XPBS for 15 min. at RT and stained with Fluoromount-G Mounting medium (with DAPI) at 1:50 000 dilution in 1XPBS for 10 min. at RT. Quantification and calculation of the autophagosome flux (*J*) and autolysosome transition state (*t_nAL_)* is performed according to an earlier publication^37^. Shortly, *J* is the calculated as the initial slope after autolysosomal induction. *T_nAL_* is calculated based on the steady-state (the number of autolysosomes before induction [v=1] is equal to the number after induction [v=2]) divided by *J*.

### Lysosomal assays and pharmacological treatments

Lysotracker Red (Invitrogen, #L7528,) was used to stain lysosomes and Lysosensor Green (Invitrogen, #L7435) to monitor the lysosomal pH on fixed cells. A working concentration of 50 nM for Lysotracker and 500 nM for Lysosensor was used. Both dyes were diluted in prewarmed conditioned culture medium (1:1 with fresh medium) corresponding to the respective cell type. After 1h of light protected incubation at 37°C/CO_2_ the cells were washed with ice cold 1XPBS and subsequently processed with immunocytochemistry. For pharmacological modulation of lysosomal function, neurons were either treated with 10 μM SB202190 (Selleck chemicals, #S1077-25mg) for 4h to enhance lysosomal activity. Alternatively, neurons were treated with 25 μM ML-SA1 (MedChemExpress, #HY-108462) either acutely for 20 min. or continuously, with treatment renewed every other day. To inhibit lysosomal function, neurons were treated with 200 nM BafA1 (Invivogen, #tlrl-baf1) for 15 min.

### Ectopic expression of CHD2 and NCOA7

The following plasmids were used to ectopic CHD2: pRP-CBh>hCHD2[NM_001271.4](ns):T2A:Puro and pRP-EF1α>hCHD2[NM_00127.4]:BGHpA-hPGK>Puro. 8×10^5^ single iPSCs were nucleofected with 5 μg of the plasmid using the P3 Primary Cell 4D-Nucleofector Kit in combination with the 4D Nucleofector Unit X. The nucleofected cells were selected with 0.25 µg/ml puromycin after 24 hours, and the next day autophagosome flux was performed. To ectopic express NCOA7, we used the following lentiviral plasmid: pLV-EF1α>hNCOA7[NM_001199619.2]>TagBFP2. The plasmid was packaged into lentiviral particles using the packaging vectors psPAX lentiviral packaging vector (Addgene #12260) and pMD2.G lentiviral packaging vector (Addgene #12259). To measure Lysosensor and autophagosome flux, 20.000 single iPS cells were transduced with the plasmid. The plasmids were constructed as a service by VectorBuilder.

### Transmission electron microscopy

Cells were fixed in 2% glutaraldehyde in 0.1M sodium cacodylate (pH 7.4) buffer for 1 hr at RT, washed in buffer and post-fixed for 1h at RT in 1% osmium tetroxide and 1% potassium ferrocyanide in 0.1 M cacodylate buffer. After being washed in buffer, cells were dehydrated in an ascending series of aqueous ethanol and were subsequently transferred via a mixture of ethanol and Epon to pure Epon as embedding medium. Ultrathin sections (80 nm) were cut, contrasted with 2% aqueous uranyl acetate, counterstained with lead citrate and examined in a JEOL JEM 1400 electron microscope operating at 60 kV.

### RNA extraction

For total RNA extraction, organoids were transferred into 1,5 ml tubes, washed twice with cold 1XPBS and snap frozen in dry ice for long-term storage at -80°C. For each RNA extraction, 3 organoids were lysed in 1000 μl of QIAzol Lysis Reagent (Qiagen, #79306). Then, 200 μl of chloroform was added per 1 ml of QIAzol Lysis Reagent used and the samples were mixed thoroughly. After a 3-4 minutes incubation, samples were centrifuged at 15’000g for 15 minutes at 4°C. The aqueous phase was transferred into a clean 1,5 ml tube and 1 μl of GlycoBlue coprecipitant (Thermo Scientific, #AM951) was added. After 5 minutes of incubation, the RNA was precipitated by adding 500 μl of isopropanol. Samples were incubated overnight at -80°C to facilitate RNA precipitation. The following day, samples were centrifuged at 15’000g for 15 minutes at 4°C. The supernatant was discarded, and RNA pellets were washed twice with 500 μl of ice-cold 75% ethanol. Finally, pellets were air-dried at RT and resuspended in the appropriate amount of Nuclease-free water (Thermo Fisher Scientific, #AM9937). Purified RNA was quantified using a Nanodrop spectrophotometer and RNA quality was checked with an Agilent 2100 Bioanalyzer using the RNA nano kit (Agilent, 5067-1512). Genomic DNA was extracted through salting out following the protocol published by McManus lab (https://mcmanuslab.ucsf.edu/protocol/dna-isolation-es-cells-96-well-plate).

### Library preparation for RNA-seq

Library preparation for RNA-sequencing was performed according to the stranded standard input TruSeq Total RNA sample preparation protocol (Illumina). cDNA library quality was assessed on an Agilent TapeStation using the High Sensitivity D1000 kit. Libraries were sequenced on a NovaSeq machine (Illunina) at a read length of 50 bp paired-end and a coverage of 20 million reads per sample.

### Whole Genome Sequencing and variant calling

Library preparation was performed with the Illumina DNA PCR-free Library Prep and Tegmentation kit (Illumina, #20041795) using 300 ng of gDNA as input. Libraries were sequenced on a NovaSeq platform using a 2×150 configuration. Variant calling was performed using DeepVar (https://doi.org/10.1038/nbt.4235) included in Nextflow (10.1038/s41587-020-0439-x) Sarek pipeline (10.1093/nargab/lqae031, 12688/f1000research.16665.2) v.3.1.1.

### RNA-seq data analysis

RNA-seq FASTQ files for organoids data were quantified at the gene level using Salmon with the seqBias, gcBias, validateMappings flags. The GRCh38 human reference (gencode v35) was used for annotation and gene-level quantification. For RNA-seq FASTQ files of iNeurons co-cultured with rat astrocytes, reads were aligned with STAR v.2.7.9a (10.1093/bioinformatics/bts635) to retain only the reads uniquely mapping to the human reference. Briefly, a combined index was generated using the human GRCh38 primary genome assembly and Rattus norvegicus 6.0 (release 104) references and gtf annotations introducing a unique identifier for the species, and reads were aligned in quantMode=GeneCounts and retaining only uniquely mapped reads. Finally, reads were disambiguated based on their mapping to the reference of either species. Differential expression analyses were performed independently for hiPSCs, neurons and organoids, comparing heterozygous or ancestralized conditions to the isogenic control. Genes with expression levels higher than 2 cpm in at least 3 samples were analyzed for differential expression. After normalization and estimation of dispersion, differential expression was tested with DESeq2 v.1.34.0 (10.1186/s13059-014-0550-8). When analyzing data from organoids, the information about differentiation rounds was used as a covariate in the statistical model. Differentially expressed genes (DEGs) were selected imposing as thresholds FDR < 0.05 and FC > 1.5. Gene ontology enrichment analyses were performed with TopGO v.2.46.0 using the ‘weight01’ algorithm and ‘fisher’ statistics. Gene set enrichment analyses were performed using the GSEA function in the clusterProfiler library. Significance for DEGs overlap was tested with the GeneOverlap R library v.1.30.0. For enrichment analyses, all genes tested for differential expression were used as universe.

### Epigenomic analysis

An atlas of putative active enhancers was generated starting from Roadmap Epigenomics data and leveraging H3K4me1, H3K27ac, H3K27me3 and H3K4me3 to define regulatory regions. We collected data from Roadmap Epigenomics for 29 histone modification profiled across H1- and H9-derived embryonic stem cells, neuronal precursors, glutamatergic neurons and primary neuronal tissues. Active enhancers were identified as intergenic and intronic regions bearing H3K4me1 and H3K27ac and devoid of H3K4me3 and H3K27me3. Intersections were performed using the *bedtools* v2.31.0 suite.

### Transcription factor binding affinity

Transcription factor binding sites prediction was performed using the TFBSTools R package v.1.31.2, leveraging motif position weight matrices (PWMs) obtained from the JASPAR database. Ancestral and contemporary alleles were analyzed to identify differential binding, focusing on sequence-specific transcription factor binding at enhancers. Motifs were matched to genomic sequences using strand-specific scoring. Confidence thresholds were set at 80%, and filtering based on p-values was applied.

### Organoids dissociation and fixation for scRNA-seq

Organoids were collected in a 24 multiwell plate, washed twice with HBSS (Stemcell Technologies, #37150) and buffers freshly prepared. Papain powder was activated in a solution composed by 1.1 mM EDTA, 0.067 mM β-mercaptoethanol (ThermoFisher Scientific, #31350-010) and 5.5 mM L-cysteine (Sigma-Aldrich, #C7880-100G) for 30 minutes at 37 °C. The correct amount of papain powder was reconstituted in the activation solution having considered the lot specific % of protein and activity. To prepare the dissociation solution, activated papain solution and DNAse I (Stemcell Technologies - #07900) were diluted in Hank’s Balanced Salt Solution (HBSS, Stemcell Technologies - #37150) to the final concentrations of 30 units/mL of papain and 125 units/mL of DNase I. Organoids were incubated for 40 minutes at 37 °C on an orbital shaker at 120 rpm and then triturated 5/6 times at RT with a p1000 to obtain a suspension consisting primarily of cell aggregates. The cell aggregates suspension was then incubated again at 37°C for 15 minutes on an orbital shaker set to 120 rpm, triturated 4/5 times at RT to obtain a suspension consisting primarily of single cells, transferred to a 15 ml tube containing 3 ml of ice-cold ovomucoid protease inhibitor solution (Worthington Biochemical, #LK003182) and centrifuged at 300 g at 4 °C for 5 minutes. Remaining cell aggregates were removed filtering the cell suspension through a 70 μm strainer (Sigma/Merk, #H13680-0070). In order to fix the samples for the Chromium fixed RNA profiling protocol (10X Genomics, # 1000414), cells were centrifuged at 200 g for 10 minutes at 4°C, the supernatant was removed, the pellet resuspended with 1 ml of fixation buffer and incubated for 20 hours at 4°C. Cells were then centrifuged at 850 g for 5 minutes at RT and the pellet resuspended with 1 ml of quenching buffer. The single cells solution was supplemented with 100 μl of pre-warmed enhancer (10x genomics, PN-2000482) and 275 μl of 50% glycerol for long-term storage at -80°C.

### Single cell RNA-seq data preprocessing

All analyses were performed within the scanpy (10.1186/s13059-017-1382-0) single cell analysis framework. Feature matrices were imported from Cell Ranger and basic filtering strategies were applied. Droplets with potential technical issues or containing more than one cells were filtered out. Only cells with less than 10% mitochondrial genes counts were retained. Cells with less than 200 detected genes and genes detected in less than 100 cells were excluded from the analysis. Cell counts were then normalised and log transformed using *sc.pp.normalize_total* and *sc.pp.log1p* scanpy functions. We regressed out the effect of total counts via *sc.pp.regress_out* and scaled the data via *sc.pp.scale* functions. All functions were run with default parameters. HVGs were detected with the *sc.pp.highly_variable_genes* scanpy function specifying the experimental replicate as batch_key, with min_mean=0.0125, max_mean=5 and min_disp=0.5. PCA was computed on defined HVGs, and the samples within each dataset were integrated via scanpy’s harmony implementation. Cells’ neighbourhood graph was computed on the top 12 or 20 principal components (pcs) with neighbourhood size (n) = 50 for the ancestralized and heterozygous organoids dataset, respectively. All scRNA-seq analyses were done using scanpy v.1.9.1, anndata v.0.9.2 and umap v.0.5.3; the matplotlib v.1.20.2 and seaborn v.0.11.1 packages were used for UMAPs and visualization of gene expression at single cell level; pandas v.1.2.4, numpy v.1.20.2, scipy 1.10.1, scikit-learn v1.1.1 statsmodels v.0.13.2 were used for data handling.

### Pseudobulk RNA-seq analysis from single cell RNA-seq experiments

To perform differential expression on scRNA-seq data and compare control lines with matched heterozygous or ancestralized lines, we generated pseudobulks. Raw reads were aggregated by cell type and replicate using adpbulk (github.com/noamteyssier/adpbulk). Genes found in at least 2 replicates with at least 5 reads were kept for differential expression analysis. Differential expression analysis was performed using edgeR as per state-of-the-art benchmarking (doi.org/10.1038/s41467-021-25960-2; doi.org/10.1093/nar/gkw448). Differential expression analysis was performed using previously published edgeR wrapper for bulk RNA-seq “edg1” (10.1126/sciadv.aaw7908), using estimateDisp function (robust=TRUE) for dispersion estimation. FDR <= 0.01 and FC >= 1.5 were used as thresholds. Gene Ontology (GO) enrichments were performed with enrichGO (10.1089/omi.2011.0118) filtering false discovery rate (FDR) at 0.05.

### Somatodendritic reconstructions

Imaging of MAP2-stained neurons was done by using Zeiss Axio Imager Z1. Somatodendritic reconstructions were performed digitally by using the NeuroLucida 360 software (Version 11, MBF-Bioscience, Williston, ND, USA). We quantified the number of dendrites, the number of dendritic ends, the soma area, and the total and mean dendritic length. To investigate the complexity of dendrites in regard to their distance from the soma, Sholl analysis was performed. We quantified the number of intersections, the number of nodes and the total dendritic length per 10 mm interval.

### Synapse quantification

Synaptic images were acquired using the Zeiss Axio Imager Z1 microscope. Regions of interest (ROIs) were traced along the proximal dendrites of neurons using MAP2 staining in ImageJ. Synaptic puncta were analyzed with the SynBot plugin^64^, using the Ilastik thresholding method to detect presynaptic Synapsin1 and postsynaptic Homer1b/c puncta, followed by co-localization analysis to detect functional synapses. Functional synaptic puncta within the traced dendrite ROIs were quantified using a custom-written ImageJ macro, calculated per 10 µm dendrite length, and averaged across dendrites for each neuron. These values were then used for statistical analysis and data visualization.

### Micro-electrode array recording and analysis

MEA recordings were performed as previously described^62,65,66^. Time-stamps per electrode were generated from Axion Biosystems raw.spk files using custom code wrote in MATLAB. Electrode bursts were detected using Maximum Spike Interval algorithm, and setting maximum spike interval to 0.05 s. Electrode bursts were restricted to be constituted by 5 spikes at least and to be longer than 0.1 ms. Consecutive electrode bursts separated by less than 0.1 s were merged. Network bursts (NB) were detected using a modified version of calculate_network_bursts function from meaRtools R package^67^.

NB intervals were identified by thresholding the combined signal with the Otsu global thresholding algorithm^68^, requiring the involvement of at least 40% of the electrodes and a duration of more than 0.1 s. In addition, we imposed a restriction on NB, requiring a minimum firing rate of 9 Hz in at least 0.25 of the actively participating electrodes.

#### Principal component and UMAP analysis

Principal component (PCA) and UMAP analysis were performed considering all activity and burst-related variables in networks from DIV28 onwards. Neuronal networks showing < 3 NB were excluded. We determined the mean value of each parameter by grouping them based on cell line and DIV. Then, PCA was performed on the averaged parameters with *prcomp* function. UMAP was performed on the first 3 PC with *umap* function, and setting random_state = 123. The distance between the averaged samples was calculated based on the two UMAP dimensions. This distance was then employed for hierarchical clustering using the *pheatmap* function from the pheatmap R package.

### Statistical analyses

All analyses performed with R were performed with R version 4.1.1 except for the analysis of transcription factor binding motifs performed using R version 4.2.2. All analyses performed with python were performed with python version 3.8.10. Statistical details related to Figure 5 can be found on https://github.com/GiuseppeTestaLab/CHD2_release.

## ACKNOWLEDGEMENTS

We thank Svante Pääbo for providing the chimpanzee and bonobo iPSCs. We thank the European School of Molecular Medicine (SEMM), where O.L. is enrolled as a PhD student in Systems Medicine, and F. Marinaro for project management support.

This work was supported by the following funding sources:

The Netherlands Organisation for Health Research and Development ZonMw grant 91217055 (to H.v.B., N.N.K., M.M., J.V. and H.J.S.).

Simons Foundation (SFARI) Grant 890042 (to N.N.K.). Leakey Foundation Research Grant 2025 (to E.L).

C. B. was supported by Project PID2023-146627NB-I00 (Spanish Ministry of Science, Innovation, and Universities CIENCIA/AEI), project 2021-SGR-313 (AGAUR/Generalitat de Catalunya) and a Leonardo Grant for Researchers and Cultural Creators, BBVA Foundation.

T.S.B was supported by the Netherlands Organisation for Scientific Research (ZonMw Vidi, grant 09150172110002).

RD was supported by a China Scholarship Council (CSC) PhD Fellowship (201906300026 to RD) for her PhD studies at the Erasmus Medical Center, Rotterdam, The Netherlands.

Work carried out in G.T.’s laboratory at Human Technopole and IEO was funded by the European Union’s Horizon 2020 research and innovation program grants RE-MEND (101057604) and R2D2-MH (101057385).

Funding bodies did not have any influence on study design, results, and data interpretation or final manuscript.

