## Supplementary data for "Human-Specific Regulation of *CHD2* Controls the Tempo of Cortical Development through Lysosomal Homeostasis"

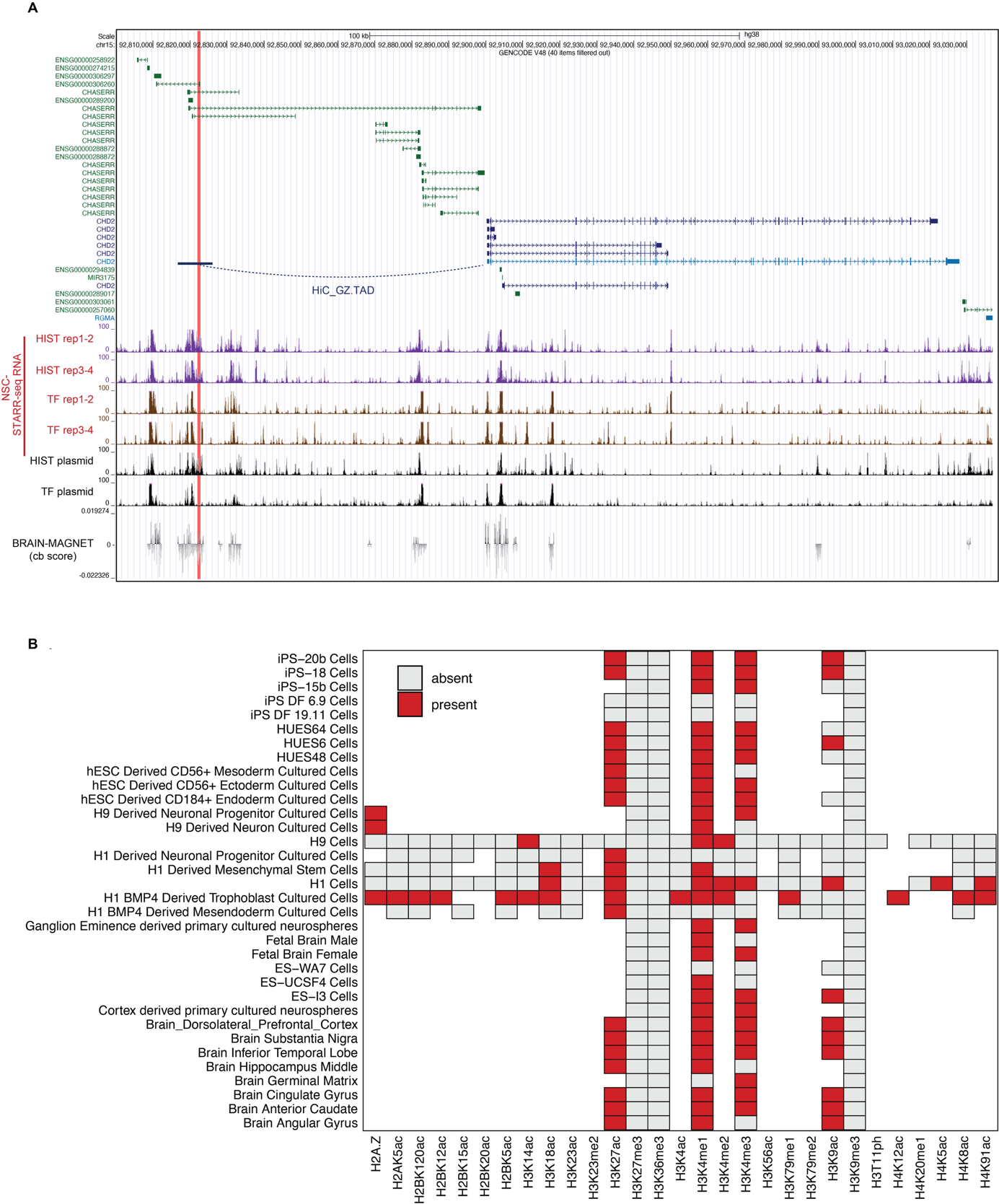

**Figure S1. The HFV-containing upstream enhancer physically contacts the CHD2 promoter in fetal cortex and is marked by active regulatory chromatin in stem cell and neural tissues.**

(A) Genome browser view of the *CHD2* locus showing the high-frequency modern-human variant (red vertical line) located within an upstream candidate enhancer. Gene tracks are shown at the top, including *CHD2* and the regulatory lncRNA CHASERR. Fetal cortical Hi-C data indicate that the HFV-containing enhancer lies within the same topological domain as *CHD2* and forms a chromatin contact with the *CHD2* promoter, supporting assignment of the enhancer to *CHD2*. Tracks below show chromatin accessibility and regulatory activity in neural stem cells from STARR-seq RNA replicates, transfection controls, plasmid input controls, and BRAIN-MAGNET nucleotide-level contribution scores across the locus.

(B) Matrix plot showing the overlap between the *CHD2* candidate regulatory region and Roadmap Epigenomics histone modification peaks across pluripotent stem cells, hESC-derived germ-layer derivatives, neural progenitor and neuronal cultures, primary neurospheres, fetal brain, and adult brain tissues. Rows indicate cell or tissue types, and columns indicate histone marks. Red squares denote histone marks detected at the regulatory region, whereas gray squares indicate absence. The region shows recurrent enrichment for enhancer- and active chromatin-associated marks, including H3K4me1, H3K27ac, and H3K9ac, across stem cell and neural contexts, supporting its annotation as an active regulatory element.

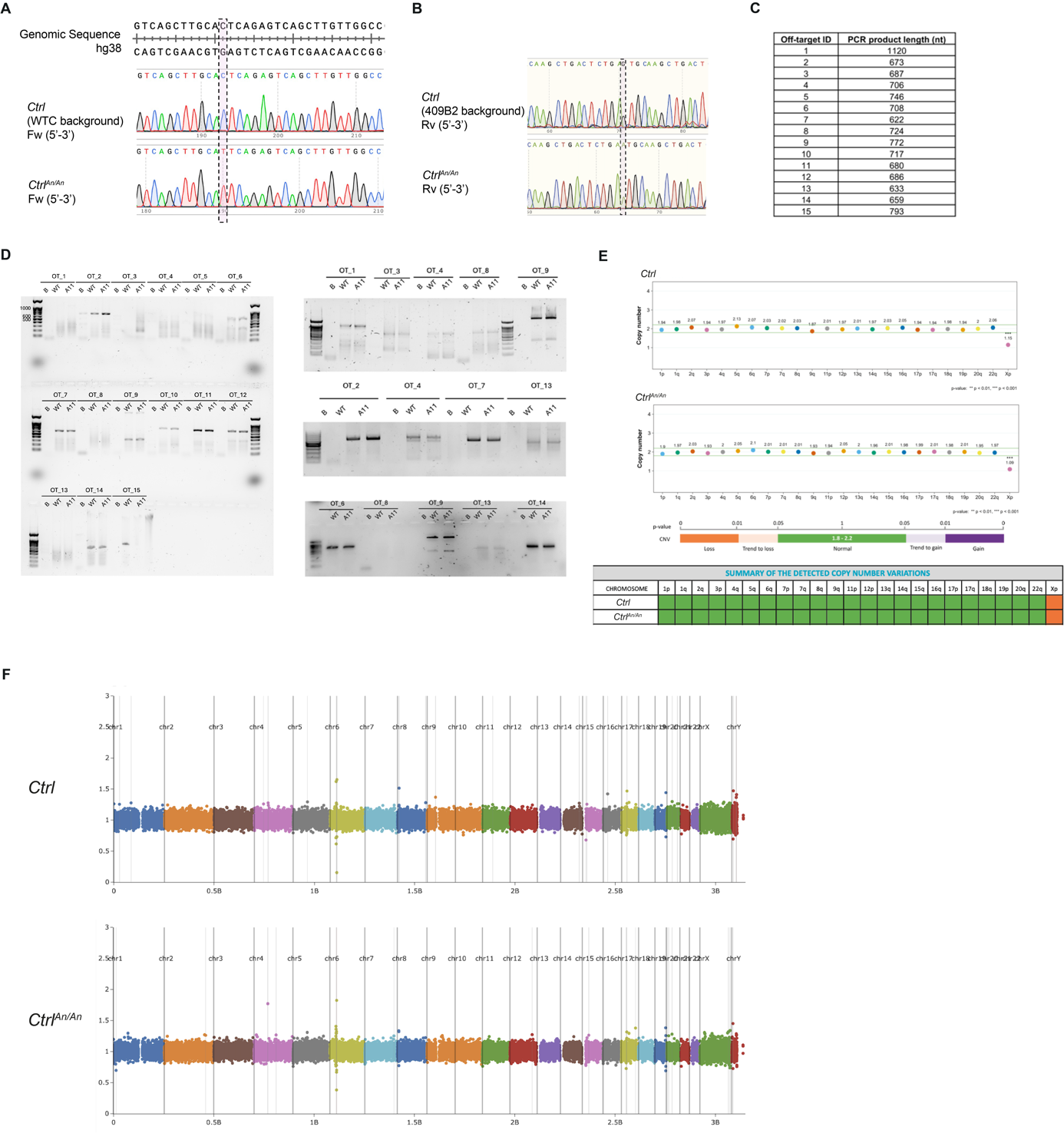

**Figure S2. *Ctrl^An/An^* CRISPR validation in *Ctrl* WTC and 409B2 genetic background**

(A-B) Sanger sequencing chromatograms confirming precise single-base editing of the target site, with wildtype (*Ctrl*) and ancestral (*Ctrl^An/^*^An^) sequences aligned to the reference genome (hg38). The dashed box highlights the edited SNV (C to T).

(C) Expected size of PCR products from 15 off-target regions with up to 3 mismatches with the single guide RNA sequence used for SNV editing.

(D) Lanes represent samples from wildtype (WT), ancestral clone (A), and blank control (B). Bands of expected sizes across all samples confirm the absence of significant off-target modifications, supporting the specificity of the CRISPR-editing process.

(E) Results of digital droplet PCR performed by Stem Genomics to detect genomic abnormalities. The orange box indicates a loss of X chromosome in both *Ctrl* (409B2) and *Ctrl^An/An^*(409B2)-derived iPSCs.

(F) Whole genome sequencing performed on *Ctrl* (WTC) and *Ctrl^An/An^* (WTC). Genome-wide scatter plots showing normalized coverage (y-axis) across chromosomes (x-axis) for whole-genome sequencing (WGS) data from *Ctrl* and *Ctrl^An/An^*. Each dot represents a genomic region, with colors indicating different chromosomes. Clustering around the baseline indicates uniform read depth. This quality control analysis validates the absence of copy number variations (CNVs) or large-scale genomic rearrangements in *Ctrl^An/An^* line and the *Ctrl* line.

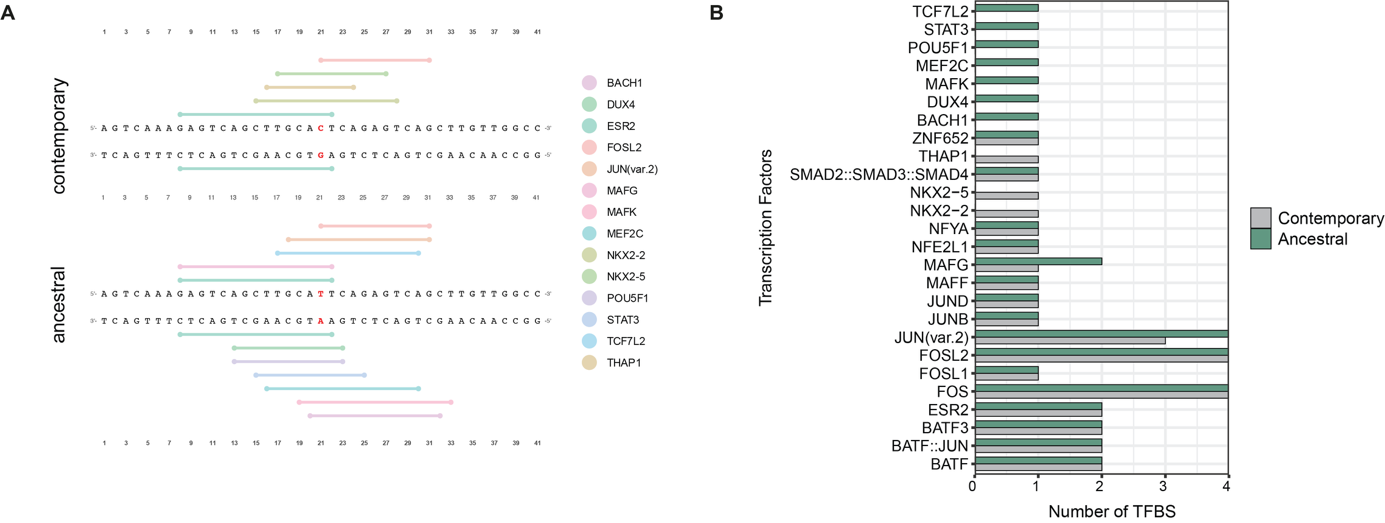

**Figure S3. The contemporary-human SNV remodels transcription factor binding site architecture at the CHD2 enhancer**

(A) Predicted transcription factor binding sites within a 41-bp sequence window centered on the *CHD2* enhancer SNV in the contemporary-human and ancestral alleles. The variant position is highlighted in red, and colored segments indicate predicted binding sites for transcription factors with high-confidence motifs in either allele.

(B) Quantification of predicted transcription factor binding sites in the contemporary-human and ancestral enhancer sequences. Bars indicate the number of predicted binding sites for each transcription factor, highlighting allele-dependent remodeling of the local regulatory landscape, including of ESR2- and MAFG-associated motifs.

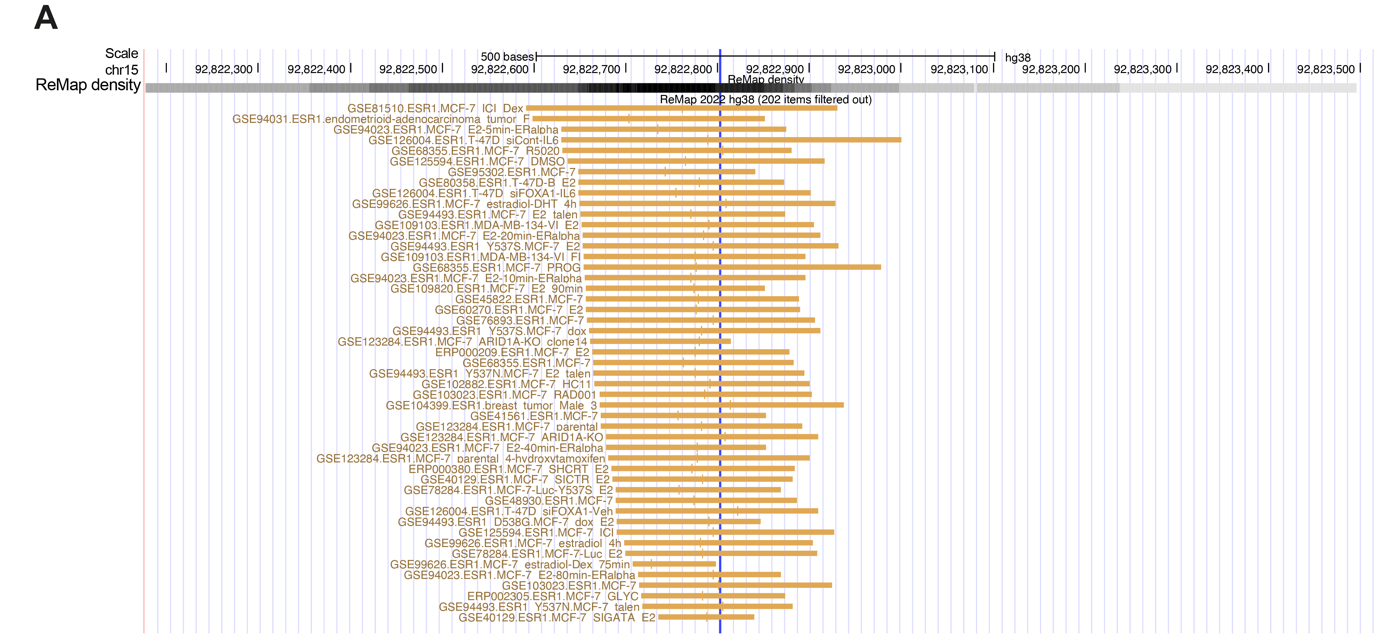

**Figure S4. ESR binds to the enhancer region in presence of the SNV.**

(A) Representative view of the peaks from publicly available ChIP-seq data included the ReMap project for ESR. The blue vertical line indicates the SNV.

Table S1. Clinical information *CHD2*-related patients used in this study

|  |  | *Patient 1* | *Patient 2* |
| --- | --- | --- | --- |
| Demographics | **Gender** | Male | Female |
|  | **Ethnicity** | Causian | Causian |
| Mutation | **gDNA coordinates** | g.93510631_93510632dup | g.92942964C>A |
|  | **cDNA coordinates** | c.2077_2078dup | c.948C>A |
|  | **Protein change** | p.Glu694* | p.Tyr316* |
|  | **Molecuar consequence** | nonsense | nonsense |
|  | **Inheritence** | *De novo* heterozygous | *De novo* heterozygous |
|  | **Clinvar** | Unknown variant | Known variant |
| Epilepsy onset | **Condition** | Developmental and epileptic encephalopathy | Developmental and epileptic encephalopathy |
|  | **Age of onset** | 18 m.o. | 7y |
|  | **Seizure type** | Tonic clonic seizures  Tonic seizures  Atypical absences  EEG shows fotosensitivity | Tonic clonic seizures  Absences  EEG shows fotosensitivity |
|  | **Trigger for first seizure** | Fever | Stress |
| Motor and cognitive development | **Before epilepsy onset** | Low Normal (76-78) | Disharmonic profile  Normal verbal IQ (91), Performal IQ below normal (74) |
|  | **After epilepsy onset** | Behavioral problems  Autistic features  Stagnation of cognitive development | Delayed Motor development (DCD 2014- scolioseis 2014)  Behavioral problems |
|  | **ID status (mild, moderate, or severe)** | Mild ID (IQ 49-60) | Mild ID (65-78) |

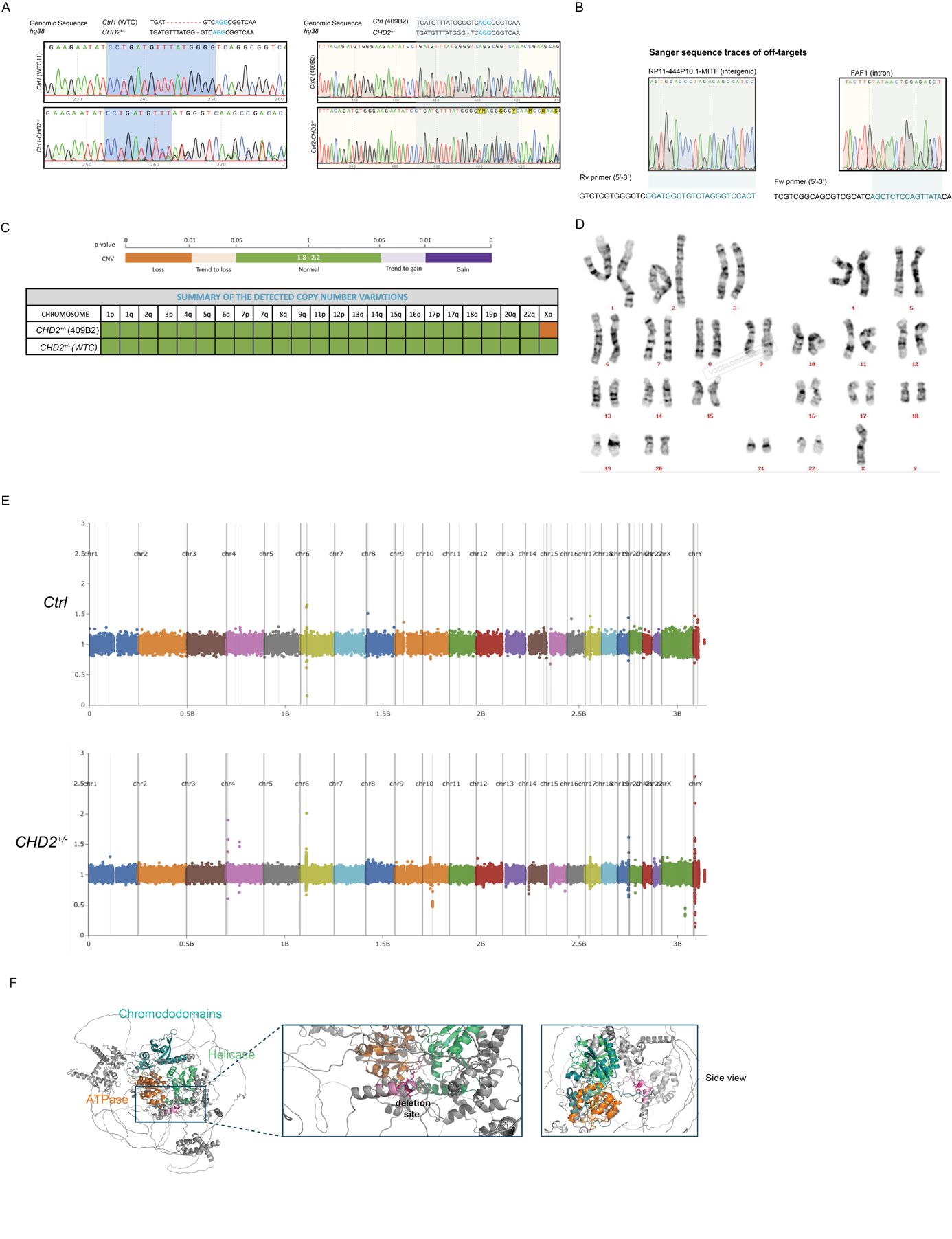
**Figure S5. *CHD2^+/-^* CRISPR validation in *Ctrl* WTC and 409B2 genetic background**

(A) Sanger sequencing chromatograms confirming *CHD2* gene editing in exon 4 in two independent controls (WTC and 409B2). The Protospacer Adjacent Motif (PAM) sequence is shown in blue. *CHD2^+/-^* (WTC) shows a bi-allelic event, with 9 nucleotide deletion in allele 1 (in-frame) and 1 nucleotide deletion in allele 2. *CHD2^+-/^* (409B2) shows a heterozygous deletion of 1 nucleotide.

(B) Off-target analysis of the two top off targets of the sgRNA used in this study. We show no genome editing in RP11-44P10.1 MITF and FAF1.

(C) Results of digital droplet PCR performed by Stem Genomics to detect genomic abnormalities. The orange box indicates a loss of X chromosome in *CHD2^+-/^* (409B2).

(D) Karyotype confirming the loss of X chromosome in *CHD2^+/-^*(409B2).

(E) Whole genome sequencing performed on *Ctrl* (WTC) and *CHD2^+/-^* (WTC). Genome-wide scatter plots showing normalized coverage (y-axis) across chromosomes (x-axis) for whole-genome sequencing (WGS) data. Each dot represents a genomic region, with colors indicating different chromosomes. Clustering around the baseline indicates uniform read depth. This quality control analysis validates the absence of copy number variations (CNVs) or large-scale genomic rearrangements in CRISPR-edited clones and the wildtype line.

(F) In addition to the intended loss-of-function mutation, a 9bp deletion was identified in the region chr15:92927265-92927273 (corresponding to exon 4 of the coding sequence of *CHD2*) on the second allele of the *CHD2^+/-^* (WTC background) cell line. The deletion is located far from functional domains both in the linear ammino acid sequence and the three-dimensional folded structure of the CHD2 protein, suggesting no impact on its structural and functional integrity.

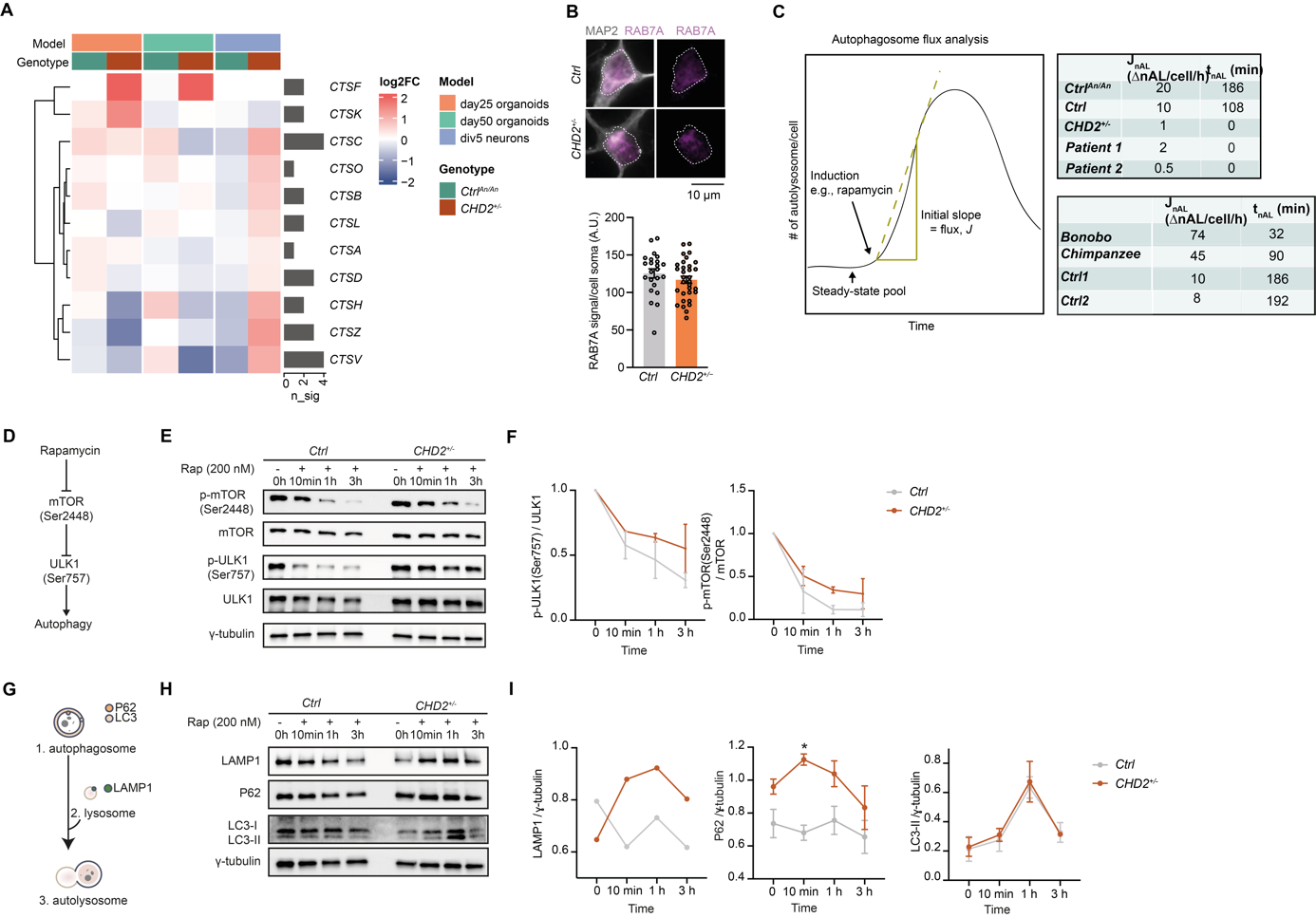

**Figure S6. Characterization of cathepsins and the autolysosomal pathway in *CHD2^+/-^* neurons**

(A) Heatmap of cathepsin-related DEGs in DIV 5 cortical neurons and Day 25/50 organoids. The heatmap shows log_2_ fold changes in *Ctrl^An/An^* and *CHD2^+/-^* neurons relative to their corresponding controls.

(B) (top) Representative images of early endosomes stained by Rab7A in *Ctrl* and *CHD2^+/-^*-derived neurons. (bottom) Quantification of RAB7A signal per soma in *Ctrl* compared to *CHD2^+/-^*-derived neurons. (n = 23 for *Ctrl* and n = 32 for *CHD2^+/-^*).

(C) (left) Schematic illustrating the autophagosome flux assay. Following induction of autophagy (e.g., rapamycin), the increase in autolysosome number over time is monitored. The initial linear rise in autolysosomes (green dashed line) is used to calculate the autophagosome flux (J), while the pre-induction autolysosome population represents the steady-state pool. (right) Table shows flux rate (*J_nAL_*) and transition state (*t_nAL_*).

(D) Induction of autolysosomal pathway by rapamycin (200 nM) leading to inhibition of mTOR by dephosphorylation of Ser2448, following inhibition of ULK1 by dephosphorylation of Ser757.

(E) Representative western blots showing reduction of phosphorylated Ser2448 site of mTOR after induction of autophagy by 200 nM rapamycin (10 min, 1h, 3h), and reduction of phosphorylated Ser751 of ULK1 in *Ctrl* and *CHD2^+/-^*-derived neurons.

(F) Quantification of phosphorylated ULK1 divided by basal ULK1, and quantification of phosphorylated mTOR divided by basal mTOR in *Ctrl1* compared to *CHD2^+/-^*-derived neurons.

(G) Schematic scheme of the autolysosomal pathway with the markers used in western blot analysis. P62 and LC3 reside in autophagosomes. LAMP1 is located in lysosomes.

(H) Representative western blots showing expression of the autophagy markers indicated in panel G in *Ctrl* and *CHD2^+/-^*-derived neurons after induction of autophagy with 200 nM rapamycin.

(I) Quantification of autophagy markers LAMP1, P62, and LC3-II in *Ctrl* and *CHD2^+/-^*-derived neurons.

Data represent means ±SEM. *p<0.05, unpaired t-test.

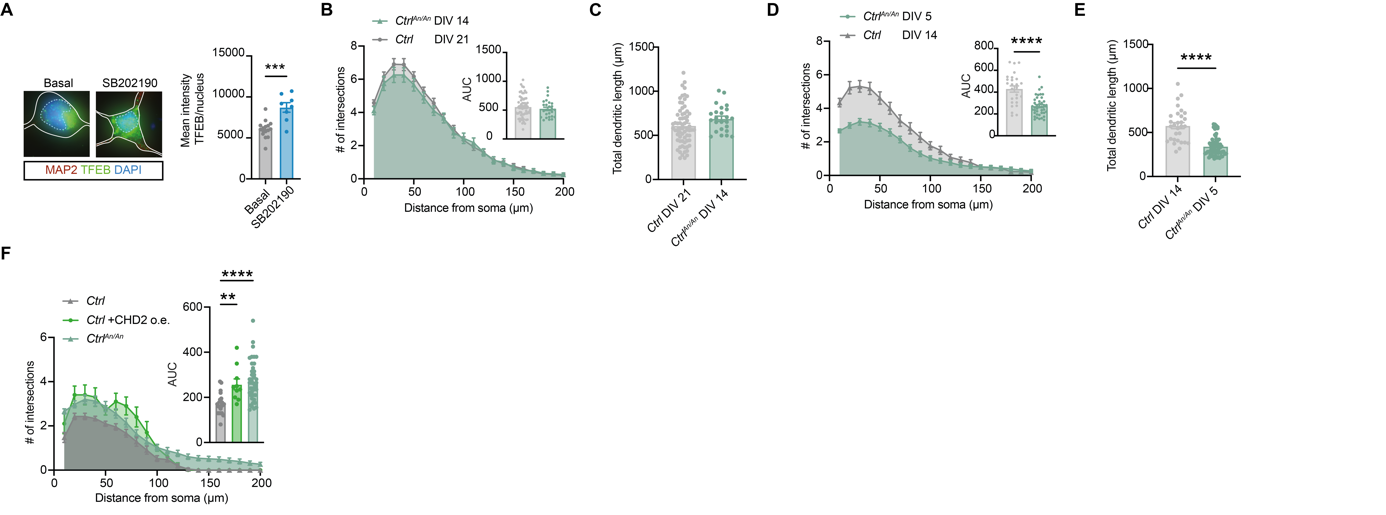

**Figure S7. TFEB translocation upon SB202190 and developmental timing comparisons between *Ctrl* and *Ctrl^An/An^* cortical neurons.**

(A) (left) Representative images of DIV 2 neurons stained for MAP2 and TFEB in *Ctrl-* and *Ctrl* neurons upon SB202190 treatment. (right) Quantification of TFEB mean intensity in the nucleus. Each dot represents the quantification of one cell; data are pooled from two independent experiments (n = 10 for *Ctrl* [WTC] and n = 10 for *Ctrl* [WTC]+SB202190).

(B) Morphometric comparison between *Ctrl* neurons at DIV 21 and *Ctrl^An/An^* at DIV 14. Quantified parameters include Sholl intersections. Insets show area under the curve (AUC) in which each dot represents one cell; data are pooled from two independent experiments (n = 30 for *Ctrl* [n = 40 for WTC, n = 18 for 409B2], n = 33 for *Ctrl^An/An^* [n = 15 for WTC, n = 15 for 409B2]).

(C) Morphometric comparison between *Ctrl* neurons at DIV 21 and *Ctrl^An/An^* at DIV 14. Quantified parameters include total dendritic length. Each dot represents the quantification of one cell; data are pooled from two independent experiments

(n = 30 for *Ctrl* [n = 40 for WTC, n = 18 for 409B2], n = 33 for *Ctrl^An/An^* [n = 15 for WTC, n = 15 for 409B2]).

(D) Morphometric comparison between *Ctrl* neurons at DIV 14 and *Ctrl^An/An^* at DIV 5. Quantified parameters include Sholl intersections. Insets show area under the curve (AUC) in which each dot represents one cell; data are pooled from two independent experiments (n = 27 for *Ctrl* [n = 15 for WTC, n = 12 for 409B2], n = 41 for *Ctrl^An/An^* [n = 30 for WTC, n = 11 for 409B2]).

(E) Morphometric comparison between *Ctrl* neurons at DIV 14 and *Ctrl^An/An^* at DIV 5. Quantified parameters include total dendritic length. Each dot represents the quantification of one cell; data are pooled from two independent experiments

(n = 27 for *Ctrl* [n = 15 for WTC, n = 12 for 409B2], n = 41 for *Ctrl^An/An^* [n = 30 for WTC, n = 11 for 409B2]).

(F) Morphometric comparison between *Ctrl-, Ctrl^An/An^*- and *Ctrl* +CHD2o.e. -derived neurons at DIV 5. Quantified parameters include Sholl intersections. Insets show area under the curve (AUC) in which each dot represents one cell; data are pooled from two independent experiments (n = 49 for *Ctrl* [n = 29 for WTC, n = 20 for 409B2], n = 41 for *Ctrl^An/An^* [n = 30 for WTC, n = 11 for 409B2], n = 23 for *Ctrl* [WTC] +CHD2o.e.).

Data represent means ±SEM. **p<0.01, ***p<0.001, ****p<0.0001, one-way ANOVA with post hoc Bonferroni correction, and unpaired student t-test.

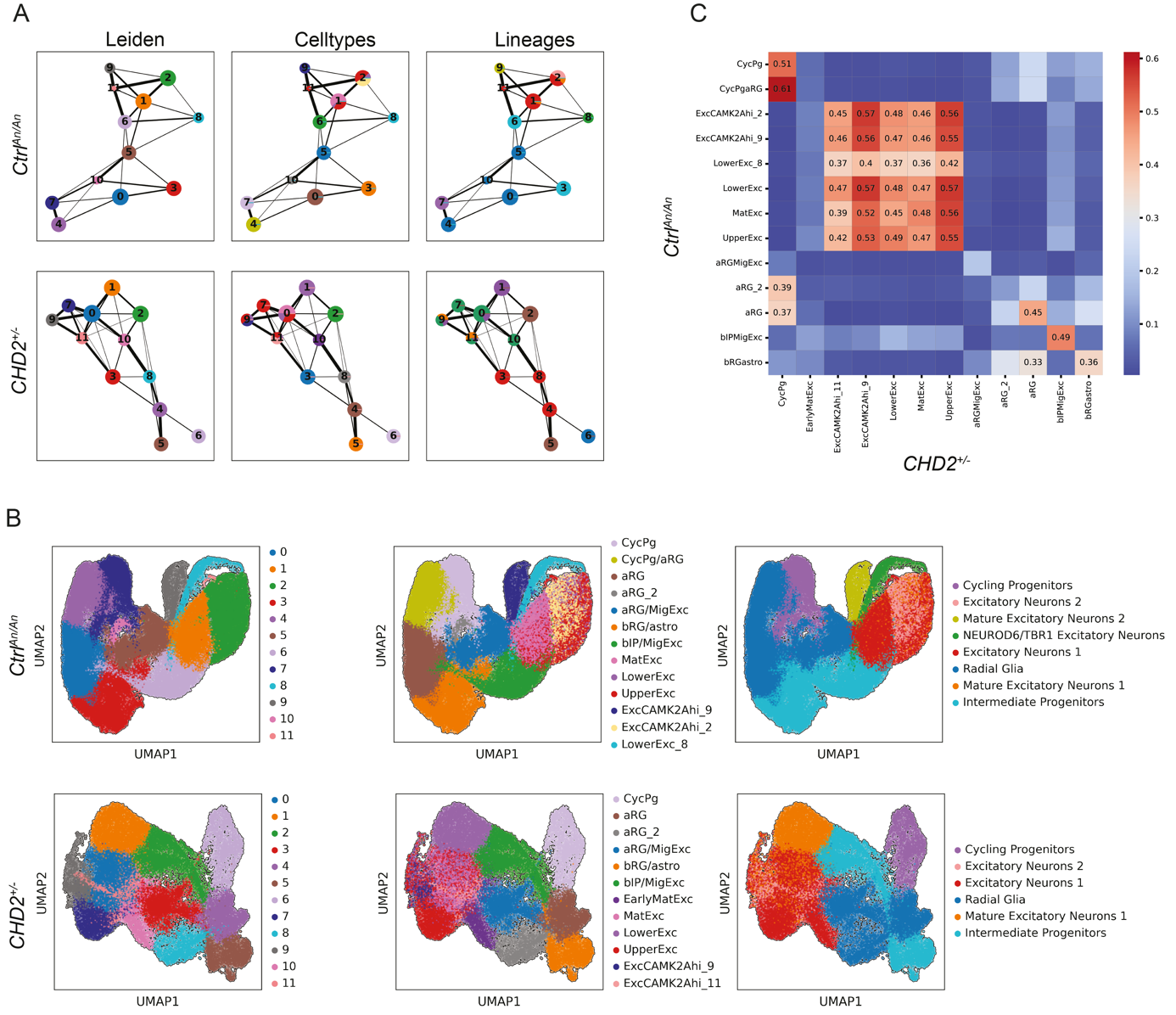

**Figure S8. Cell types and lineages annotation**

(A) PAGA graphs are respectively reported for *Ctl^An/An^* (top) and *CHD2^+/-^* (bottom) illustrating connectivity between Leiden clusters, cell types, and inferred lineages, highlighting distinct developmentally relevant connections between cell states on a graph-like structure.

(B) UMAP projections representing the transcriptome of *Ctl^An/An^* (top) and *CHD2^+/-^* (bottom) colored by clusters and their annotations (Leiden, cell types and lineages, from left to right).

(C) Heatmap of Jaccard similarity index measured for each couple of annotated cell types, based on shared genes among the top 1 thousand markers of each cell type. Here, the two scRNA-seq datasets (*Ctrl* and *Ctrl^An/An^* on the y-axis and *Ctrl* and *CHD2^+/-^* on the x-axis) show a general similarity among neuronal subtypes and a stricking similarity by cell population for all progenitors (CycPg, aRG, bRGastro) and for migratory neurons (MigExc).

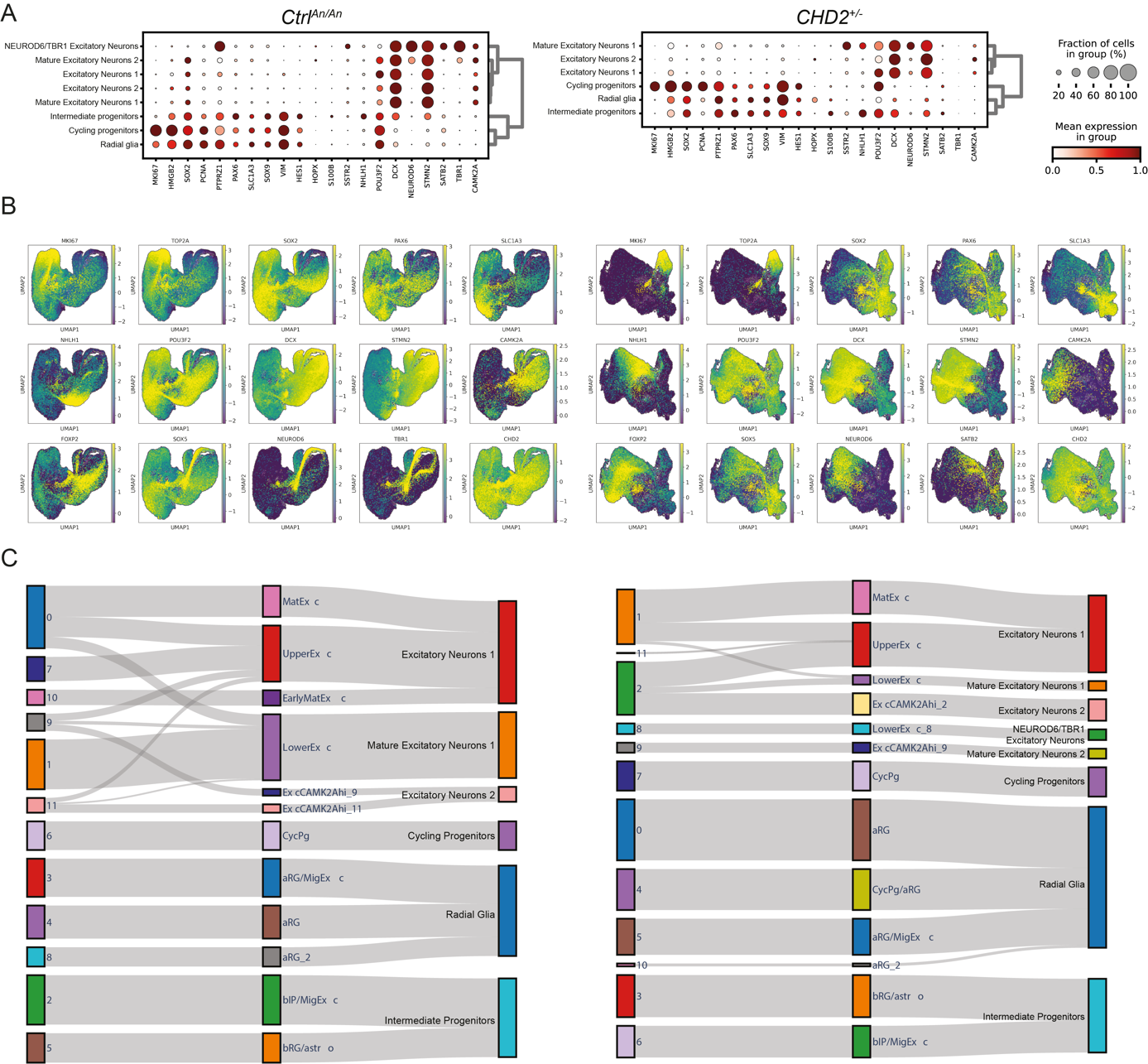

**Figure S9. Fate markers and lineage grouping.**

(A) Dot plots showing the expression of fate marker genes across annotated lineages for *Ctrl^An/An^* (left) and *CHD2^+/-^* (right) organoids. Dot size represents the percentage of cells expressing the gene within each lineage, while color intensity indicates the mean expression level.

(B) UMAP projections for *Ctrl^An/An^* (left) and *CHD2^+/-^* (right) organoids, depicting expression patterns for selected genes. Each panel corresponds to a specific gene, with color intensity reflecting expression levels.

(C) Sankey plots illustrating the relationships between Leiden clusters (left bars), annotated cell types (central bars) and lineages (right bars) for *Ctrl^An/An^* (left) and *CHD2^+/-^* organoids.

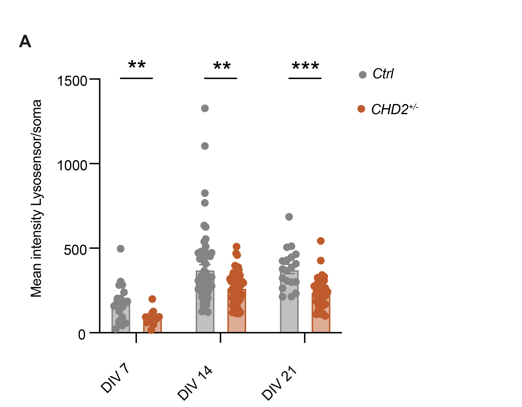

**Figure S10. Quantification of lysosomal acidification over time**

(A) Quantification of Lysosensor green mean intensity in *Ctrl* and *CHD2*^+/-^ cortical neurons at DIV7, DIV14, and DIV21.

Data represent means ±SEM. **p<0.01, ***p<0.001, one-way ANOVA with post hoc Bonferroni correction
